# Neural dynamics of stroboscopic stimulation at different stimulation frequencies

**DOI:** 10.1101/2021.06.26.450044

**Authors:** Ram Kumar Pari

## Abstract

Stroboscopic stimulation has been previously shown to induce visual hallucinations and altered states of consciousness, state by entraining the brain to the driving frequency, similar to those reported during the psychedelics although little systematic research exists on the effect of specific stimulation frequency on experience. The present study investigated the effects of different stroboscopic stimulation frequencies on neural dynamics, such as signal diversity (Lempel-Ziv complexity) and spectral power and attempted to relate these changes to self-reported changes in experiential content. The results indicated that the stimulation frequencies near the alpha band (8 – 12 Hz) caused the greatest increase across all neural measures, with 8 Hz consistently displaying the most pronounced results, relative to baseline. All tested frequencies led to an increase in all experiential dimensions, relative to baseline.

---

Consciousness or rather the idea of consciousness is a multi-faceted phenomenon that can be described as the state or the process by which the brain can process information while being self-aware of its existence and place within that information chain. In the search for a better understanding of consciousness and its neural markers, there are two popular approaches that have been used, specifically access consciousness and phenomenological consciousness. Access consciousness relates to attention, information access and the level of consciousness, such as waking vivid conscious awareness, sleep or even coma; while phenomenological consciousness focuses on the experiential content of consciousness, such as seeing the colour red or touching something hot or feeling emotions (Block, 1995, 2007; A. Seth, 2010). One thing to note with these reductionistic approaches to consciousness is that while they might functionally compartmentalise and quantise the features of consciousness, they might not represent all the individual facets that make up conscious experience (A. K. Seth, Izhikevich, Reeke, & Edelman, 2006).

Our perceptual experiences can be referred to as conscious content or contents of consciousness. Some studies have indicated that the phenomenological diversity of conscious content may relate to levels of consciousness (access and attention) with some recent theories suggesting that consciousness is better understood as a multidimensional space of information with such different levels, best understood as regions of information within this space (Boly et al., 2013; A. K. Seth, Dienes, Cleeremans, Overgaard, & Pessoa, 2008). Within this multidimensional space, levels of consciousness may act as barriers or rather gates to the information present within them (Bayne, Hohwy, & Owen, 2016; Bayne, Seth, & Massimini, 2020; Nani et al., 2019; Schwartzman et al., 2019). The idea of gating is evident when comparing states such as a normal waking consciousness versus a sleep state. During normal waking consciousness we have access to a wide range of information and have relatively diverse experiences. When it comes to more unique states such as in the case of seizures and mild anaesthetics, the range of information and diversity of experience appears much more gated compared to a normal waking state (Hohwy, 2012; Kiefer, 2008). Thereby the perceptual experience of consciousness seems to vary between these different states or levels of consciousness.

Neuromodulation is a process by which underlying brain activity can be affected or modified, as a result of some external stimuli, such as electromagnetic or chemical stimulation. Neuromodulation can be targeted specifically to a small region of the brain or even act on a system-wide basis and has been widely used in research and medical interventions and relies on the fundamental principle of being able to alter aberrant brain activity towards a more desirable form (Deer et al., 2014). Similarly, pharmacological perturbation, such as psychedelic and hallucinatory drugs have also been used to study consciousness as they induce altered states of consciousness, which are unique and specific to the drug (Riba, Anderer, Jané, Saletu, & Barbanoj, 2004). Psychedelic altered states of consciousness (ASC) have powerful phenomenological effects, such as the expansion of the visual field and its content, alterations of time and space and even changes in the perception of the self and emotions (Swanson, 2018). Given the pronounced experiential changes these pharmacological agents can cause, it seems likely that the gating for conscious content is affected, which is supported by reports of an increased range of conscious content during psychedelic states (M. M. Schartner, Carhart-Harris, Barrett, Seth, & Muthukumaraswamy, 2017). In terms of neural dynamics, psychedelics have been shown to modulate the spectral power profile, leading to a pronounced drop in alpha power (8-12 Hz) (Robin L. Carhart-Harris et al., 2016; Muthukumaraswamy et al., 2013; Riba et al., 2004), which has been classified by some as one of the neural markers of the associated visual hallucinations (Swanson, 2018).

Several attempts have been made to quantify the phenomenological change in experience with alterations in gating in conscious content that accompanies such altered states of consciousness. The approaches have ranged from scales that measure the level of consciousness through the gating of information (Directed Transfer Function Measures and Glasgow Coma scale) (Bjørn E. Juel, Romundstad, Kolstad, Storm, & Larsson, 2018; Teasdale & Jennett, 1974), to representing conscious information (Φ and Perturbation complexity Index) (Casali et al., 2013; Tononi, Boly, Massimini, & Koch, 2016) and the entropy / phenomenological diversity of conscious content (Lempel-Ziv Complexity) (Robin Lester Carhart-Harris et al., 2014). In the case of direct entropy and phenomenological diversity measures of conscious content, there appears to be a correlative link between phenomenological diversity and neural signal diversity or Lempel-Ziv complexity (Michael M Schartner et al., 2017). Lempel-Ziv complexity has been shown to be effective in detecting different states of consciousness, when applied to neural time-series, being able to differentiate between anaesthetics, regular sleep and NREM sleep and even differing psychedelic states (Farnes, Juel, Nilsen, Romundstad, & Storm, 2019; Bjørn Erik Juel, 2019; Michael Schartner et al., 2015; Michael M. Schartner et al., 2017).

A recent study by Schartner et al. (2017) reported a notable increase in Lempel-Ziv complexity across the brain during altered states of consciousness (ASC) induced by psychedelics, such as LSD, Ketamine and psilocybin, in comparison to the placebo condition. The study also showed fluctuations in the spectral power profile with changes across beta, theta and delta bands respective to each psychedelic and an overall decrease across the alpha band across all psychedelics. Behaviourally, the study used the Altered States of Consciousness Questionnaire (ASCQ) to measure the subjective experience of the psychedelic session and showed significant interactions between signal diversity and phenomenological dimensions such as ego dissolution etc. Apart from psychedelics, levels of consciousness have also been studied using sleep states such as NREM sleep and Lucid dreaming states and even using methods of neuromodulation such as stroboscopic stimulation, which uses stroboscopic light to induces simple visual hallucinations, which bears some similarities to the psychedelic state (Wackermann, Pütz, & Allefeld, 2008).

Historically, stroboscopic stimulation has been used as a recreational tool by the public through the use of machines, such as the dream machine and other self-made approaches through the last century based on a book “The Living Brain” by Walter Grey (1953) (Ter Meulen, Tavy, & Jacobs, 2009). Stroboscopically induced visual hallucinations (SIVH), the main subjective effect of such stimulation was initially discovered by Jan Purkinje in 1819 (Paul C. Bressloff, Cowan, Golubitsky, Thomas, & Wiener, 2002; 1819). SIVH’s that occur with eyes closed are described as visually intense experiences that display magnificent colours, geometric form constants and complex imagery (Bartossek, Kemmerer, & Schmidt, 2020; Paul C. Bressloff et al., 2002; Mauro, Raffone, & VanRullen, 2015; Schwartzman et al., 2019). These form constants have also been reported in various ASC such as those induced by Marijuana (Siegel & West, 1975) LSD (Knoll, Kugler, Höfer, & Lawder, 1963) models suggesting that form constants may arise out of the same neural processes and correlates (Rule, Stoffregen, & Ermentrout, 2011). Several computational models indicate the role of the early visual system, such as V1 and V2 regions, in producing the form constants observed in stroboscopic hallucinations, particularly their physical hyper column structures and how the two regions interact and synchronize with each other, with some models even indicating the physical structure of the retinal ganglion cells (Billock & Tsou, 2012; Paul C Bressloff, Cowan, Golubitsky, Thomas, & Wiener, 2001; Muthukumaraswamy et al., 2013).

Stroboscopic stimulation has also been shown to affect the spectral power profile in a similar manner to psychedelics, with results showing a consistent drop in the alpha band (8 – 12 Hz) (Bartossek et al., 2020), and even alpha power suppression after the stimulation has ceased (Streicher, 2020). A recent paper by Schwartzman et al. (2019) showed that stroboscopic stimulation can produce an ASC similar to a psychedelic induced ASC. The study found increased signal diversity across multiple stimulation frequencies and also significant shifts within the spectral power profile, which were in line to those found during psychedelic ASC (Michael M Schartner et al., 2017). The study used 3 Hz and 10 Hz stimulation conditions and a dark condition (with eyes closed) for the baseline, with each condition lasting 10 minutes. Using a modified version of the ASCQ questionnaire the study reported increases in several experiential dimensions, such as seeing patterns or strange things, muddled thinking, a more vivid imagination and even ego dissolution, which were similar to Psilocybin induced ASC. Spectrally, the study also showed an increase in beta and theta power density and a decrease in alpha power density. The study also noted the strong qualitative intensity of such stroboscopically induced experiences, such as seeing geometric configurations and visuals, flying through space, falling into the water and recollecting faces from the past of family and friends, which were associated with fine detail.

While previous studies using stroboscopic stimulation have found similar results to those of altered states of consciousness created by pharmacological interventions, a central point which has not been addressed is how different frequencies, particularly those outside the alpha band range (8 – 12 Hz) affect the underlying altered states of consciousness. Additionally, there may be a simple linear relationship between stimulation frequency and changes in measures of neural dynamics, such as signal diversity that is, as stimulation frequency increases so do measures of signal diversity.

The purpose of this study was to compare how different stimulation frequencies affect signal diversity, spectral power profile and subjective experiences. Based on previous research, seven stimulation frequencies were chosen from each spectral band. The chosen frequencies were 3 Hz, 7Hz, 8Hz, 10Hz, 13 Hz, 15 Hz and 19 Hz and the study also included a baseline dark condition. These eight conditions were then compared for significant differences in signal diversity (Lempel-Ziv), absolute spectral power, normalised spectral power (density) in relation to the baseline condition. Further, changes in neural dynamics were related to experiential changes using responses to a modified version of the Altered States of Consciousness Questionnaire (ASCQ) (see Figure 4A). The ASCQ has been previously used and validated in the research of altered states of consciousness in psychedelic and stroboscopic stimulation research (Dittrich, 1998; Michael M. Schartner et al., 2017; Schwartzman et al., 2019).

Based on past research in stroboscopic stimulation, we expect a linear relationship between stimulation frequency and changes in measures of neural dynamics. Exploratory correlation analyses were also performed across the relative difference of each condition from baseline for signal diversity, absolute spectral power, normalised spectral power (density) and each dimension of the modified ASCQ to identify if there were any links between these different measures.

## Methods

### Participants

In total, twenty-three participants took part in the experiment. At the time of the experiment, the participants were students of the University of Sussex, were healthy and above the age of 18. The participants provided informed consent and were reimbursed with ten-course credits as compensation for their time. Due to the potential risk associated with stroboscopic stimulation, participants were required to complete questionnaires on anxiety and epilepsy before taking part in the experiment. They were excluded if they scored above the exclusion thresholds (One of the participants met these exclusion criteria and was excluded). The epilepsy screening questionnaire used was based on Placencia et al. (1992), and the anxiety screening questionnaire used the State-Trait Anxiety Index, Trait Version (STAI) (Spielberger, 1983) with their exclusion criteria’s being a single positive response for the epilepsy questionnaire and above 60 for the anxiety questionnaire. Participants 9 and 11 were excluded due to either too many bad EEG channels or noisy EEG data, which was assessed visually following artefact correction, and three was removed due to missing behavioural data. Certain participants (1,2,6,13,17) were removed as outliers based on standard methods of removing outliers +/− 2 SD from the mean. Given the relatively low number of participants, the number of outliers for removal was capped at five, leaving 14 participants in total.

### Ethical Approval

This study was reviewed and approved by the University of Sussex Ethics committee (Application id ER/RP440/1).

### Procedure

The participants were seated in an electromagnetically shielded, dark and noise-proof room, 50 cm away from and facing the Lucia N°03 hypnagogic stroboscope (Innsbruck, Austria) (Fig 1). The stroboscopic light stimulation was presented at 80% of the maximum capacity of the LED output for all stimulation conditions resulting in a maximum luminous flux of 5600lm (lumen) or luminance of 6880 lux at the point where the participant’s eyes were relative to the lamp. In total there were eight conditions, each lasting for four minutes, one baseline ‘Dark’ condition (no stimulation) and seven stimulation conditions each at a different frequency, ‘3 Hz’ (strobe at 3Hz), ‘7 Hz’ (strobe at 7Hz), ‘8 Hz’ (strobe at 8Hz), ‘10 Hz’ (strobe at 10Hz), ‘13 Hz’ (strobe at 13Hz), ‘15 Hz’ (strobe at 15Hz) and ‘19 Hz’ (strobe at 19Hz). The baseline condition was always presented first, while the other stimulation conditions were presented in a randomised order. Due to the high levels of brightness of the lights and for maximal SIVH, the participants were instructed to keep their eyes closed during all conditions.

**Figure 1:**
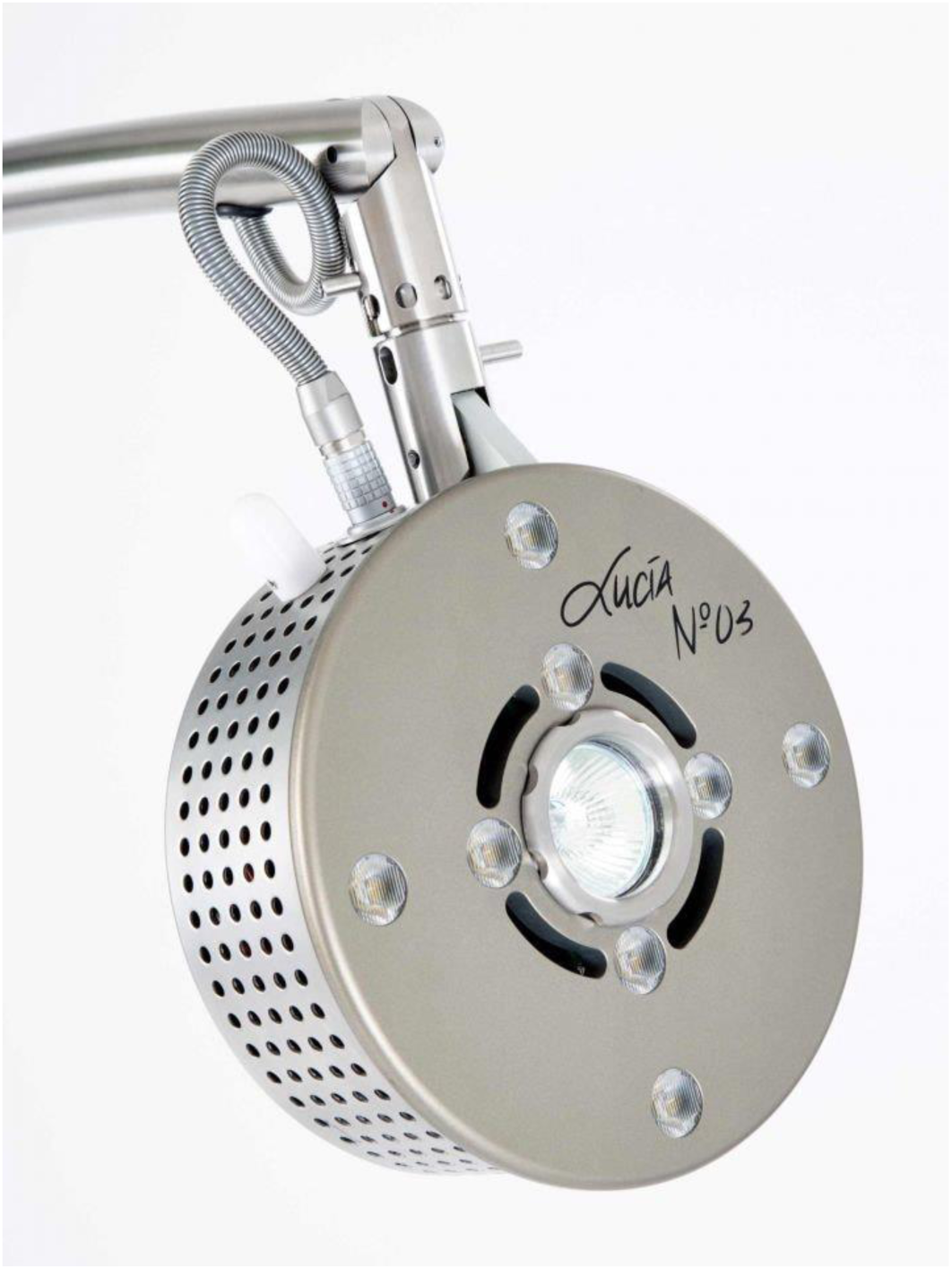
Lucia N°03 hypnagogic stroboscope (Innsbruck, Austria). The stroboscope consists of eight LEDs, a central halogen lamp and can produce stroboscopic frequencies in the range of 0.004 to 200 Hz. The present study used the LED array at a luminous flux of 5600 lumens with the participant seated 50 cm away with their eyes closed. Figure reused from Mountanalogue.co.uk (2020)

The experiment began with a practise session wherein the participants were familiarised with stroboscopic stimulation, and the questionnaire measures used in the main experiment. The practise session contained a dark condition and a stimulation condition (10Hz). Following each condition, the participant completed an abbreviated version of the Altered States of Consciousness Questionnaire (ASCQ) (Dittrich, 1998) via a computer that measured different dimensions such as Imagination, Awe, Body, Colours, Afraid, Surroundings, Float, Thought, End, Scene, z, Patterns, Lights, Drowsy and Absorbed (Figure 4A). During the practice session, the participants were encouraged to ask questions about the measures if anything was unclear or misunderstood. After this, the participants were shown an instructional video about the experiment while setting up the EEG cap and apparatus.

The main experiment consisted of the eight conditions one baseline ‘Dark’ condition (no stimulation) and seven stimulation conditions, ‘3 Hz’ (strobe at 3Hz), ‘7 Hz’ (strobe at 7Hz), ‘8 Hz’ (strobe at 8Hz), ‘10 Hz’ (strobe at 10Hz), ‘13 Hz’ (strobe at 13Hz), ‘15 Hz’ (strobe at 15Hz) and ‘19 Hz’ (strobe at 19Hz) with the stimulation conditions presented in random order and with each condition lasting for four minutes each. After the condition was over, the participants were asked to rate the average intensity of their experience across the entire condition on a scale of 0 – 100 with 100 indicating the most intense experience they’ve had, and then fill out the ASCQ questionnaire.

### Data

The EEG data was recorded using ASA-Lab 4.7.11 (ANT Neuro, Enschede) with 64-channel ANT waveguard caps, using standard Ag/AgCl electrodes and placed according to the standard international 10 - 20 system with an average reference, and using a 64-channel ANT Neuro amplifier with a sampling rate of 2048 Hz and did not use any analog filters during online recording. Impedance values of every individual electrode were kept under 40 kΩ, with electrodes in the occipital region kept at under 20 kΩ. Eye electrodes placed vertically and horizontally around the participant’s eyes were used to measure any eye movements during the study.

The data was pre-processed using custom scripts in MATLAB R2020a (Mathworks, Inc. Natick, MA, USA), with certain scripts from the EEGLAB toolbox (Delorme & Makeig, 2004), with ICLabel (Pion-Tonachini, Kreutz-Delgado, & Makeig, 2019)and ERPLAB plugin(Lopez-Calderon & Luck, 2014) and the data analyses were performed using custom MATLAB, Python and R scripts. The EEG data was downsampled to 1024 Hz, and then FIR filtered using a 1 Hz high pass filter and a 70 Hz low pass filter and any 50 Hz electrical noise was removed using CleanLine. Channels contaminated with artefacts were identified using EEGLAB based on the kurtosis of the recorded electrodes, and any electrodes with a z score of >5 were removed and interpolated using spherical interpolation from nearby electrodes. The data was then sinusoidally detrended using a custom MATLAB script to remove the steady-state visual response to the stroboscopic stimulation

### Analyses

#### Signal diversity measures

Single-channel Lempel-Ziv complexity (LZs) was used to calculate the signal diversity of the EEG data over channel Oz, using the same method from Schartner (2017) paper (github.com/mschart/SignalDiversity). The channel OZ was selected for LZs as it was previously used as the best indicator of stroboscopic stimulation and SIVH. For a given segment of data, Lempel-Ziv complexity (LZs) quantifies the signal diversity of the signal by calculating the number of distinct patterns of activity present within that segment of data and works by binarization of the multidimensional time series (see Figure 2).

**Figure 2:**
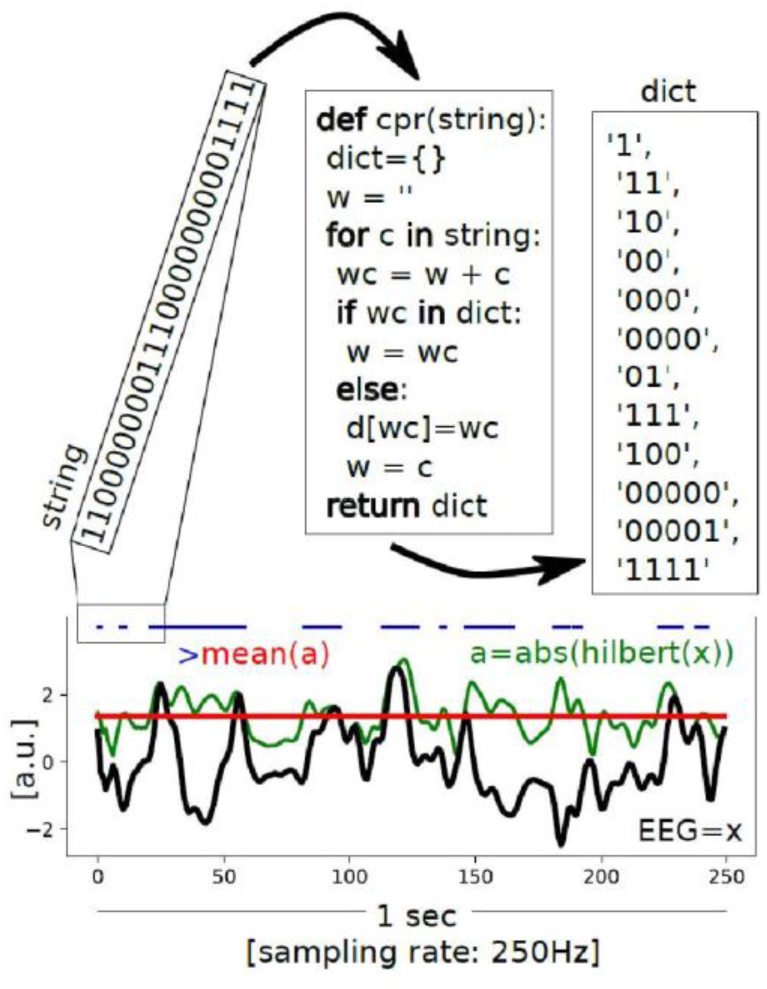
Schematic representation of the algorithm used for Lempel-Ziv calculation. The figure showcases one second, 250 Hz sampled EEG signal represented in black (x). The mean (red) of the analytic signal (green) is used to calculate the binarized signal (blue), which is then encoded into a dictionary of unique subsequences and then normalized by dividing the raw value by the randomly shuffled binary sequence. Figure from Schwartzman et al. (2019)

In short, for each condition, signal diversity of each channel in this study is calculated by dividing 4 min recordings into 10 seconds, non-overlapping segments, subtracting the mean amplitude from each of these segments diving them by standard deviation and then removing any linear trends with the SciPy signal detrend function. The continuous signal data of each channel is then converted with a Hilbert transform, using the mean of its absolute value as a threshold, into a binary string. The LZ algorithm then divides the sequence into non-overlapping and unique binary subsequences. The number of subsequences produced is proportional to the Lempel-Ziv complexity, that is, the higher the degree of randomness, the greater the number of unique subsequences it should contain and hence a higher LZs. The LZs score for each segment is then normalised by diving it with its raw value with the value obtained for the same binary input sequence that is randomly shuffled. The LZs scores range 0 to 1, with 0 representing a flat, constant signal and 1 representing a unique signal with no repeated sequences at all. Figure 2 shows a schematic representation of the algorithm used for the Lempel-Ziv complexity function. The LZs scores averaged over each condition were then compared across subjects between baseline-condition and between different conditions using paired t-tests with Bonferroni correction for multiple comparisons.

#### Altered States of Consciousness (ASCQ) analysis

The abbreviated version of the ASCQ questionnaire was used to investigate the subjective experience of each condition as a whole and to compare the level of behavioural effects that each condition had. The ASCQ questionnaire had 16 dimensions, of which four dimensions (Afraid, Thought, End and Drowsy) were classed as negative, and the other 12 were positive. A Total measure was also calculated by adding the scores of all the other ASCQ dimensions and normalizing them across the number of dimensions in the ASCQ. Due to a data collection error, Block 7 was lost across all participants. A MANOVA was used to compare if there was any change in scores across each dimension between different conditions. A post hoc Dunnett modified Tukey-Kramer Pairwise Multiple Comparison test (T3 procedure) was also conducted across each significant dimension of the ASCQ to investigate between-condition differences since paired t-tests could not be used as a result of the data loss (Dunnett, 1980).

### Spectral power profile

#### Absolute spectral power

The spectral power on of channel Oz across different power bands, Delta (1-4Hz), Theta (4-8Hz), Alpha (8-12Hz) and Beta (13-30 Hz) was calculated for each condition. Based on past research using stroboscopic stimulation, Channel OZ was selected as it has been previously indicated to be reflective of the SIVH (Schwartzman et al., 2019). For each condition, 4-minute recordings were used to calculate the mean spectral power, by decomposing the EEG signal data over channel OZ into time and frequency components, for a certain spectral band across channel OZ and repeated for each participant (absolute power). The changes across the absolute spectral power for each frequency band between conditions were analysed using paired t-tests with Bonferroni correction for multiple comparisons.

#### Normalized Spectral power

The spectral power distribution across different power bands, Delta (1-4Hz), Theta (4-8Hz), Alpha (8-12Hz) and Beta (13-30 Hz) was also calculated for each condition in order to compare the measure with other psychedelic and stroboscopic stimulation studies and to reveal changes that are not showcased by absolute spectral power. Similar to LZs, for each condition 4-minute recordings were divided into 10 second, non-overlapping segments and each of these segments (for every channel) the spectral density across each frequency band was calculated using discrete fast Fourier transform (github.com/mschart/SignalDiversity). This data was then normalized with the sum of the power of the four bands to create each frequency band represented as a percentage of the total spectral power profile (relative power). The data was then further averaged across each segment, channel and participant. The changes across the spectral density for each frequency band between conditions were analysed using paired t-tests with Bonferroni correction for multiple comparisons.

#### Correlation analysis between signal diversity, power spectra and spectral density and ASCQ

The relative differences across LZs, normalized spectral power, absolute spectral power and each dimension of the ASCQ was used to compute a Pearson correlation (r) across participants for each condition. Given the number of variables involved for such a small number of participants, the correlations were not for FWE but instead had their minimum significance level set to *p < 0.01*.

## Results

### Signal diversity

To compare changes in Lempel-Ziv scores (LZs) across conditions, paired t-tests were run with Bonferroni correction, which revealed a significant increase in LZs across all stimulation conditions 3 Hz (M = 0.468, t(13)= −3.925, *p_bon_* = .049), 7 Hz (M = 0.474, t(13)= −4.030, *p_bon_* = .040), 8Hz (M = 0.486, t(13)= −4.495, *p_bon_* = .017), 10Hz (M = 0.476, t(13)= −4.461, *p_bon_* = .018) and 19 Hz (M = 0.472, t(13)= −4.274, *p_bon_* = .025) respective to the baseline dark condition (M = 0.436). There were no significant between-condition comparisons for conditions 13 Hz (M = 0.464) and 15 Hz (M = 0.464). Fig 3 shows the distribution of LZ scores, as well as the individual scores of each participant across the different conditions. In addition, Lempel-Ziv complexity across all channels was also calculated for comparison with other studies using LZc scores (see Appendix A) (Michael M Schartner et al., 2017). The results revealed the greatest increase in LZc at 8 Hz with a general increase. In general, stimulation frequencies within or close to the alpha band (8 - 12Hz) seemed to produce the largest increases in LZs.

**Fig 3:**
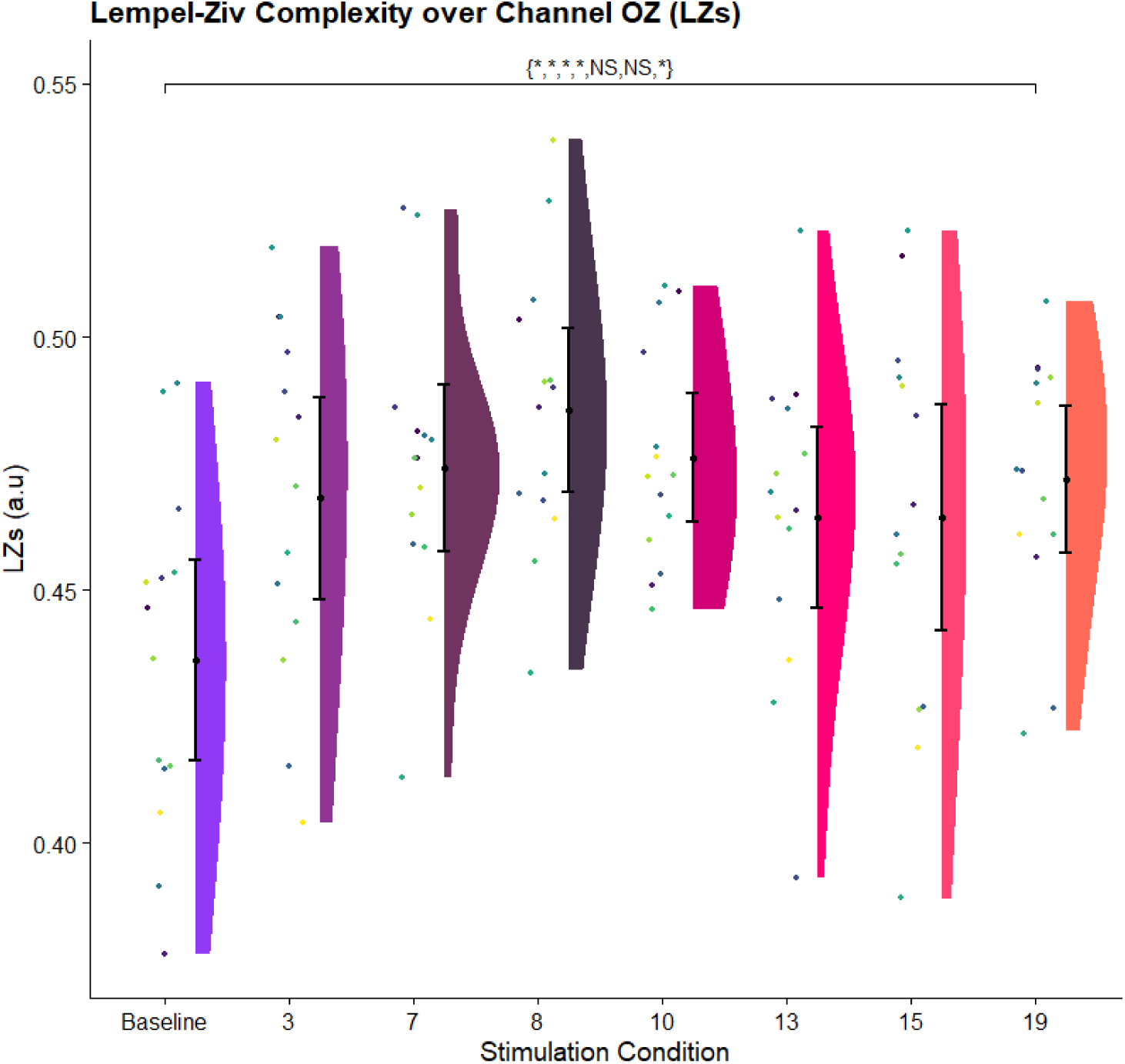
Mean Lempel-Ziv complexity scores (LZs) for channel Oz. Rain cloud plot displays the Lempel-Ziv complexity scores over channel Oz (LZs), averaged over each condition’s 4-minute window, and all participants. All comparisons are Bonferroni corrected, and asterisks denote significant comparisons, NS for non-significant, * indicates *p < 0.05*. The violin plots represent the distribution for each condition. The individual coloured dots represent individual participants results with each participant represented by a unique colour. The error bars represent the 95% confidence intervals of the data for each condition.

### Altered States of Consciousness (ASCQ) analysis

The different dimensions and average responses to the abbreviated ASCQ scale are shown for all conditions in Fig 4. The average responses across all positive dimensions displayed an increase compared with the dark baseline condition, which is in line with past stroboscopic stimulation and psychedelic studies (Robin L. Carhart-Harris et al., 2016; Muthukumaraswamy et al., 2013; Riba et al., 2004; Michael M Schartner et al., 2017; Schwartzman et al., 2019).

**Figure 4:**
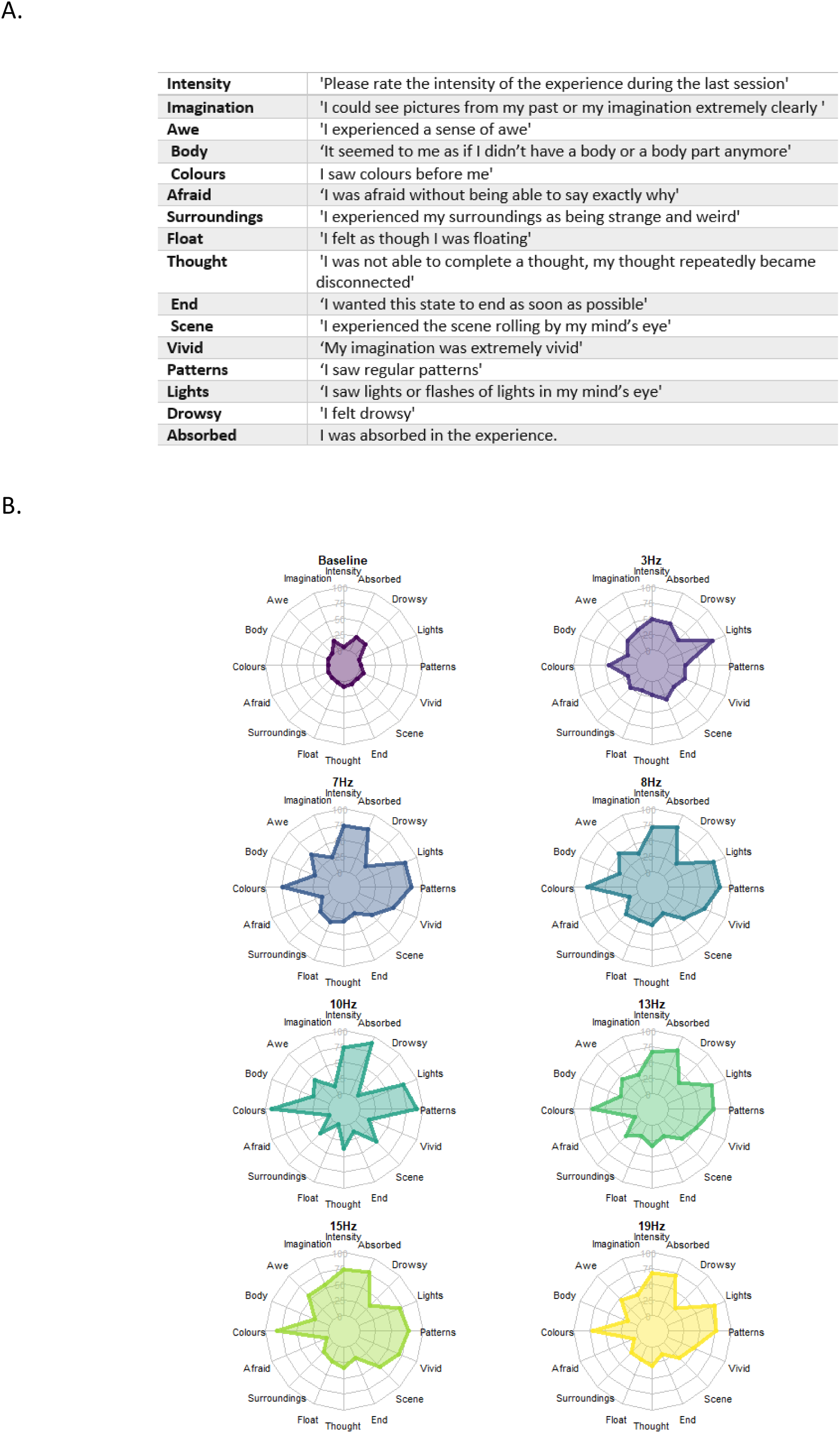
A. The individual dimensions of the abbreviated version of the ASCQ and its respective questions used in the post-stimulation questionnaire. B. Radar plots of ASCQ responses across all conditions. Plot showcases the average (across subjects) values of each ASCQ dimension across each condition represented in an individual radar plot for each condition with values from 0 to 100, with increments of 25 per concentric ring.

A MANOVA consisting of the different ASCQ dimensions as the dependent variables and conditions as the independent variable revealed a significant difference between conditions (Pillai’s Trace = 1.715, *F* = 1.926, *df* = (112), *p* = < .001). The univariate result revealed a significant difference between conditions for the dimensions Intensity (*F* = 14.633, *df* = (7), *p* = < .001), Awe (*F* = 4.193, *df* = (7), *p* = < .001), Colours (*F* = 13.944, *df* = (7), *p* = < .001), Surroundings (*F* = 2.321, *df* = (7), *p* = .030), Float (*F* = 2.259, *df* = (7), *p* = .033), Scene (*F* = 4.098, *df* = (7), *p* = < .001), Vivid (*F* = 8.803, *df* = (7), *p* = < .001), Patterns (*F* = 22.649, *df* = (7), *p* = < .001),Lights (*F* = 20.026, *df* = (7), *p* = < .001), Absorbed (*F* = 9.089, *df* = (7), *p* = < .001) and the dimensions Imagination (*F* = 2.0255, *df* = (7), *p* = .059), Body (*F* = 1.889, *df* = (7), *p* = .078) and Float (*F* = 1.975, *df* = (7), *p* = .065) was found to be close to significance.

Dunnett modified Tukey-Kramer (DTK) Pairwise Multiple Comparison tests were then conducted for post-hoc testing across ASCQ dimensions that reached significance or were close to the minimum significance level (*p* = 0.05) in the MANOVA. These results are visualized in figure 5, which showcases the 95% confidence intervals for each pairwise comparison between conditions, with significant comparisons highlighted in red. The DTK test was chosen since paired t-tests could not be run as a result of a data loss of one block (Block 7) for each participant, resulting in an unequal number of participants between conditions (unequal variance) (Dunnett, 1980).

**Figure 5:**
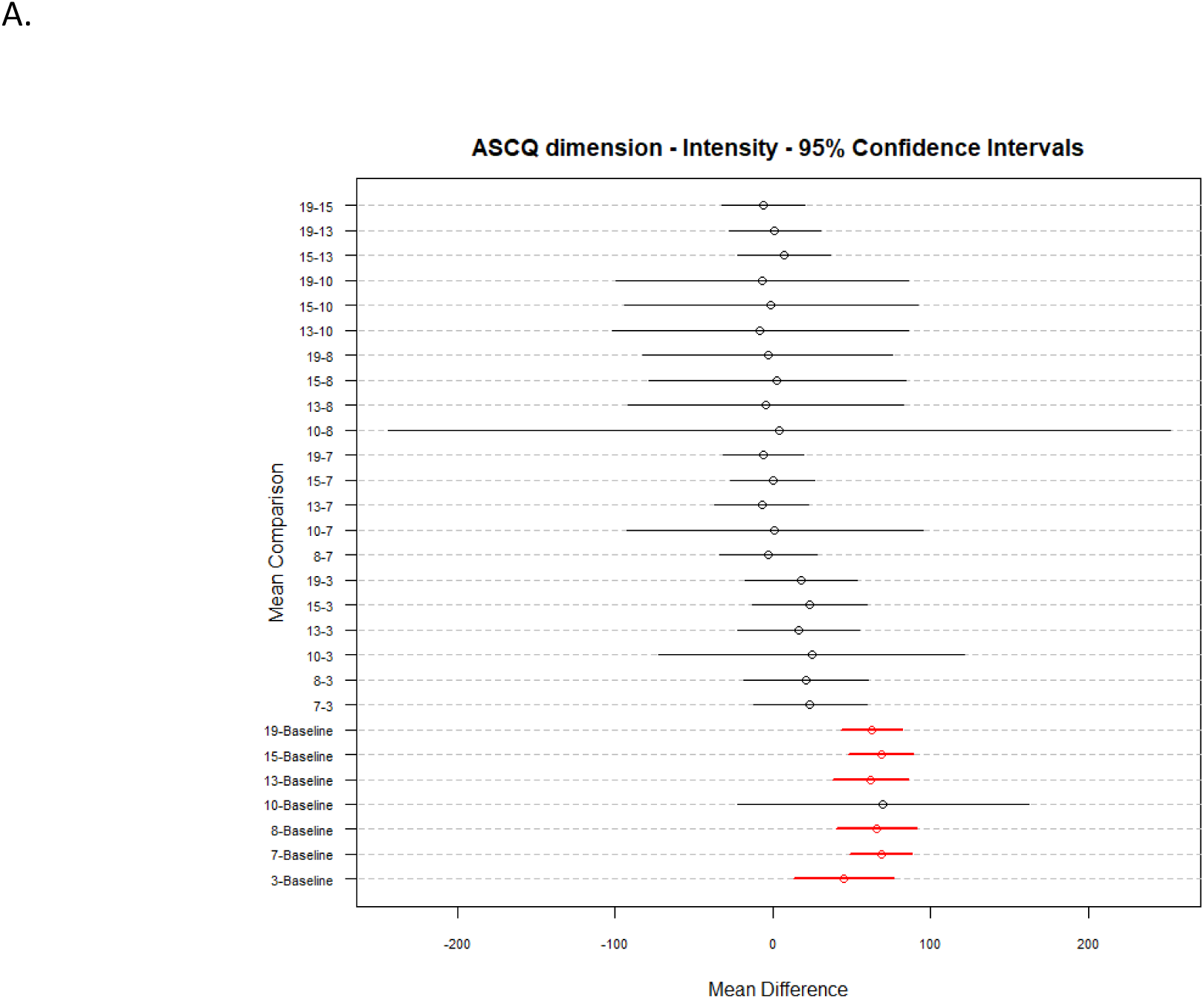
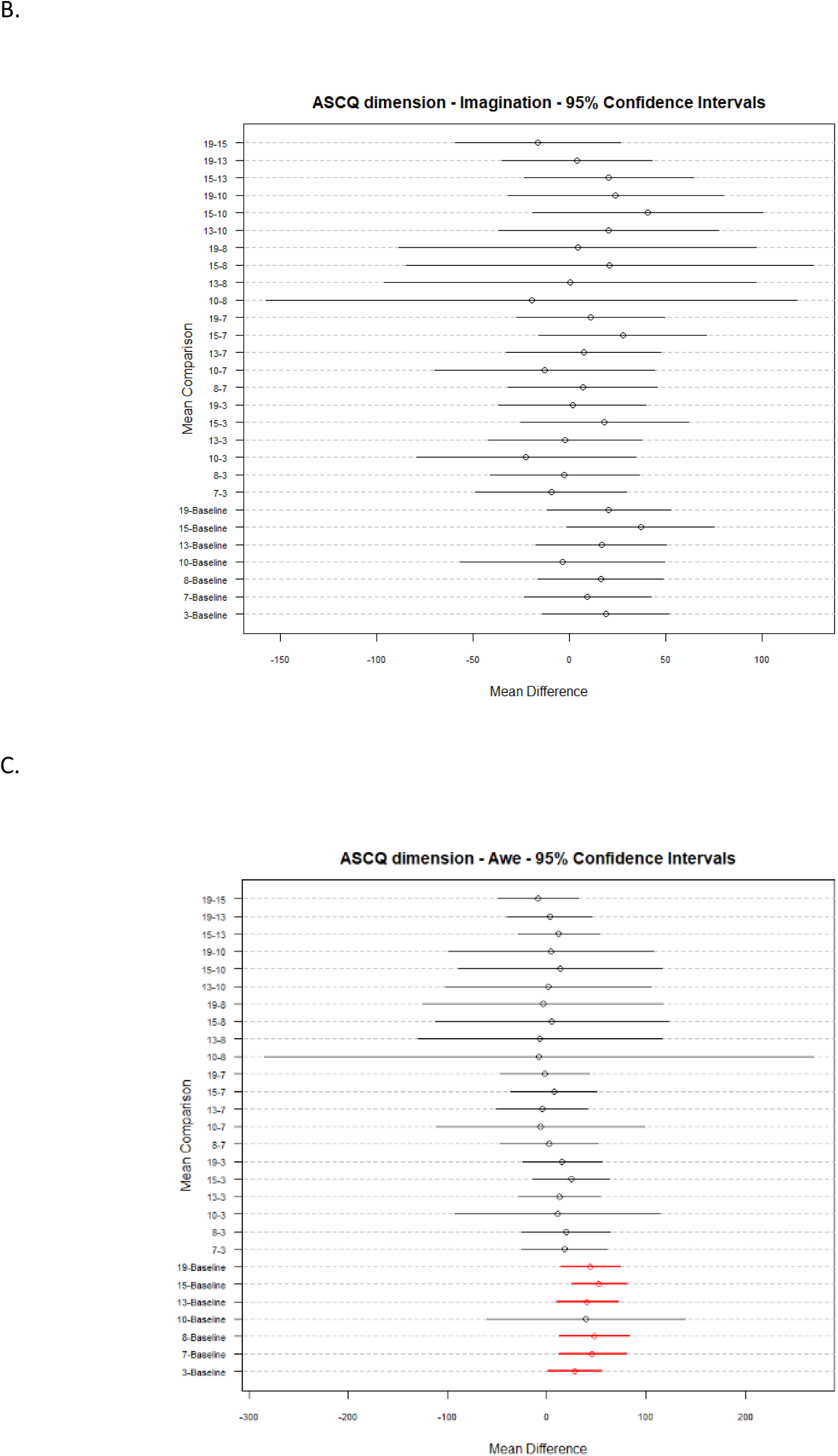
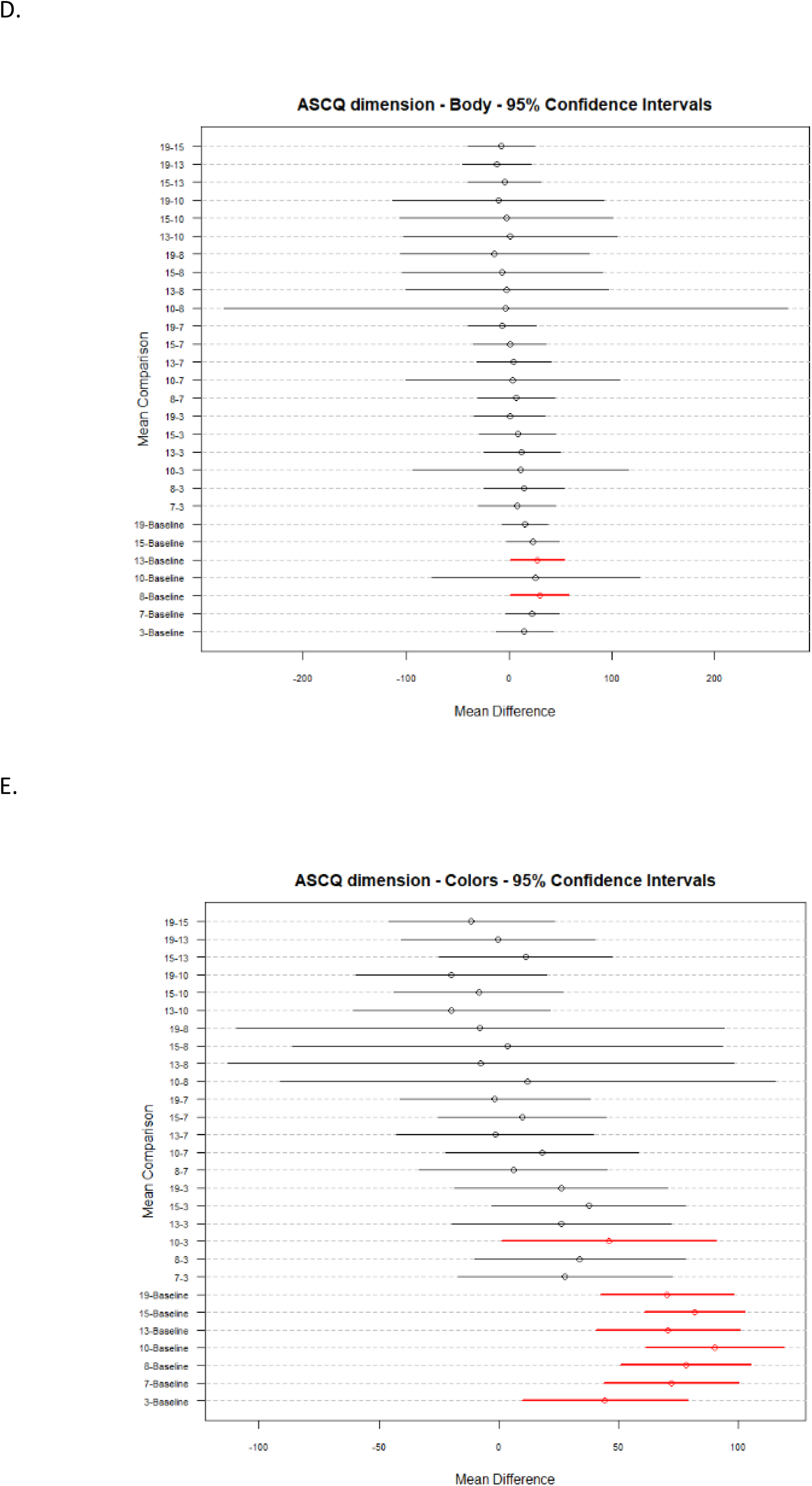
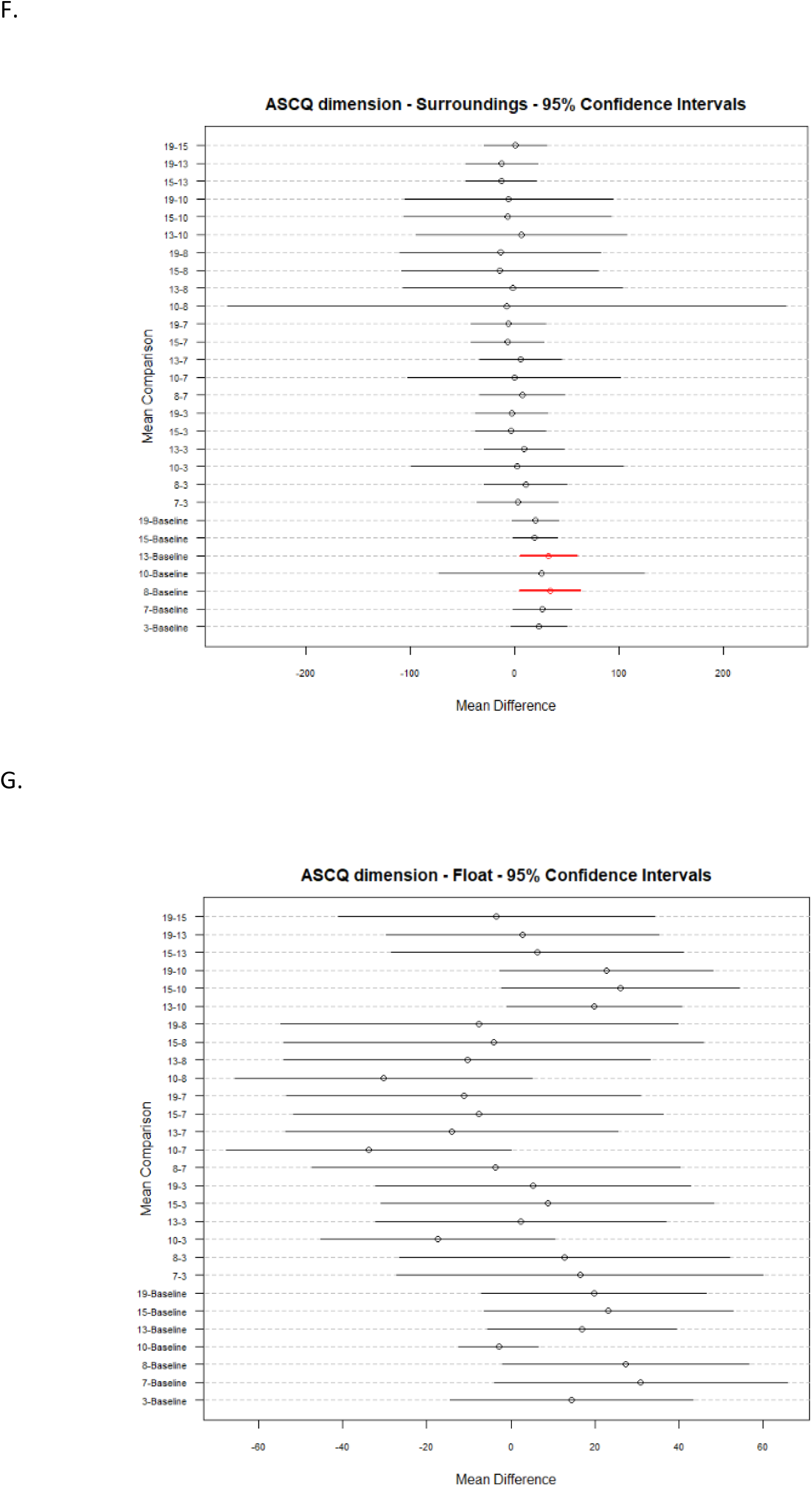
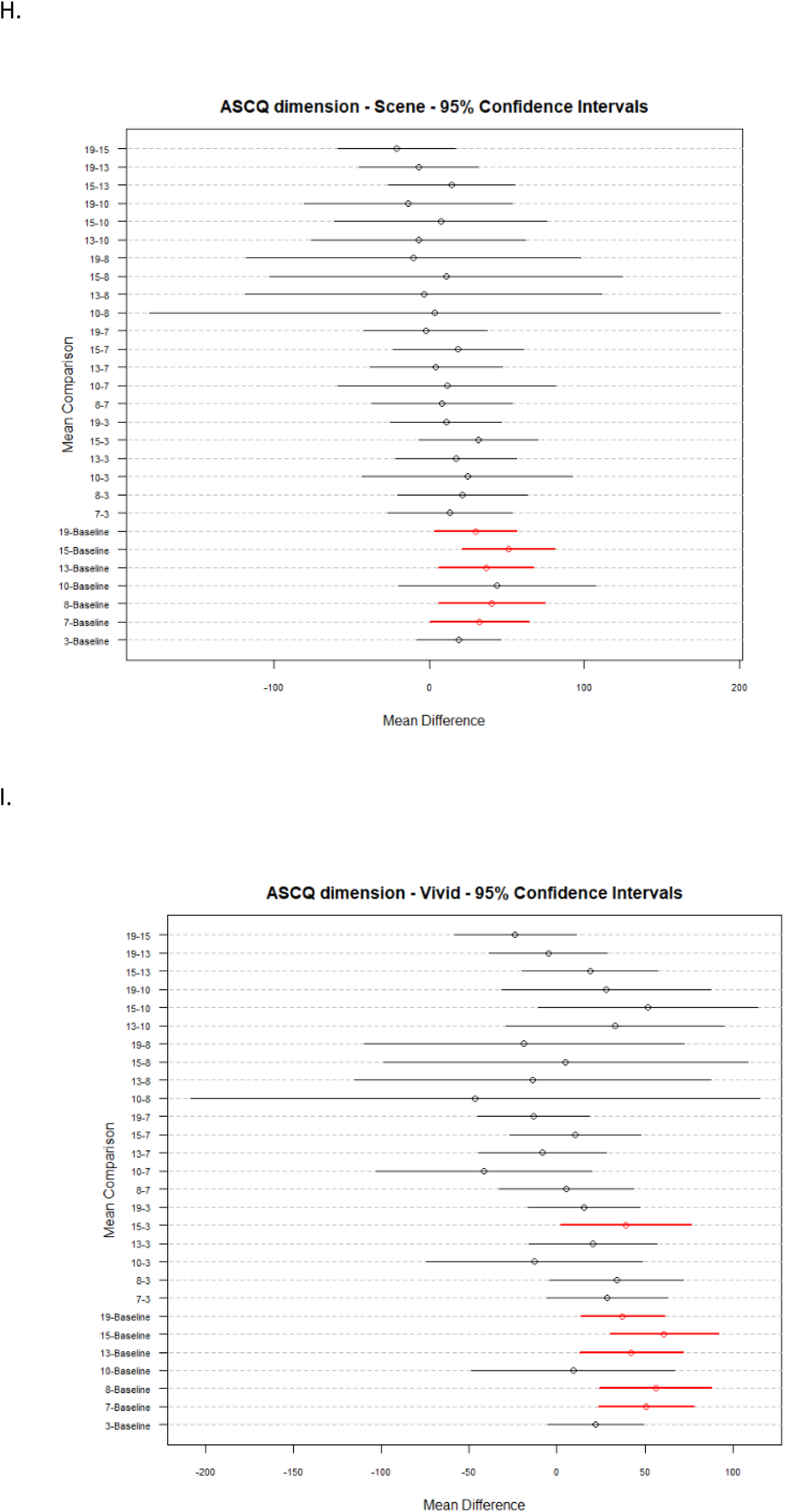
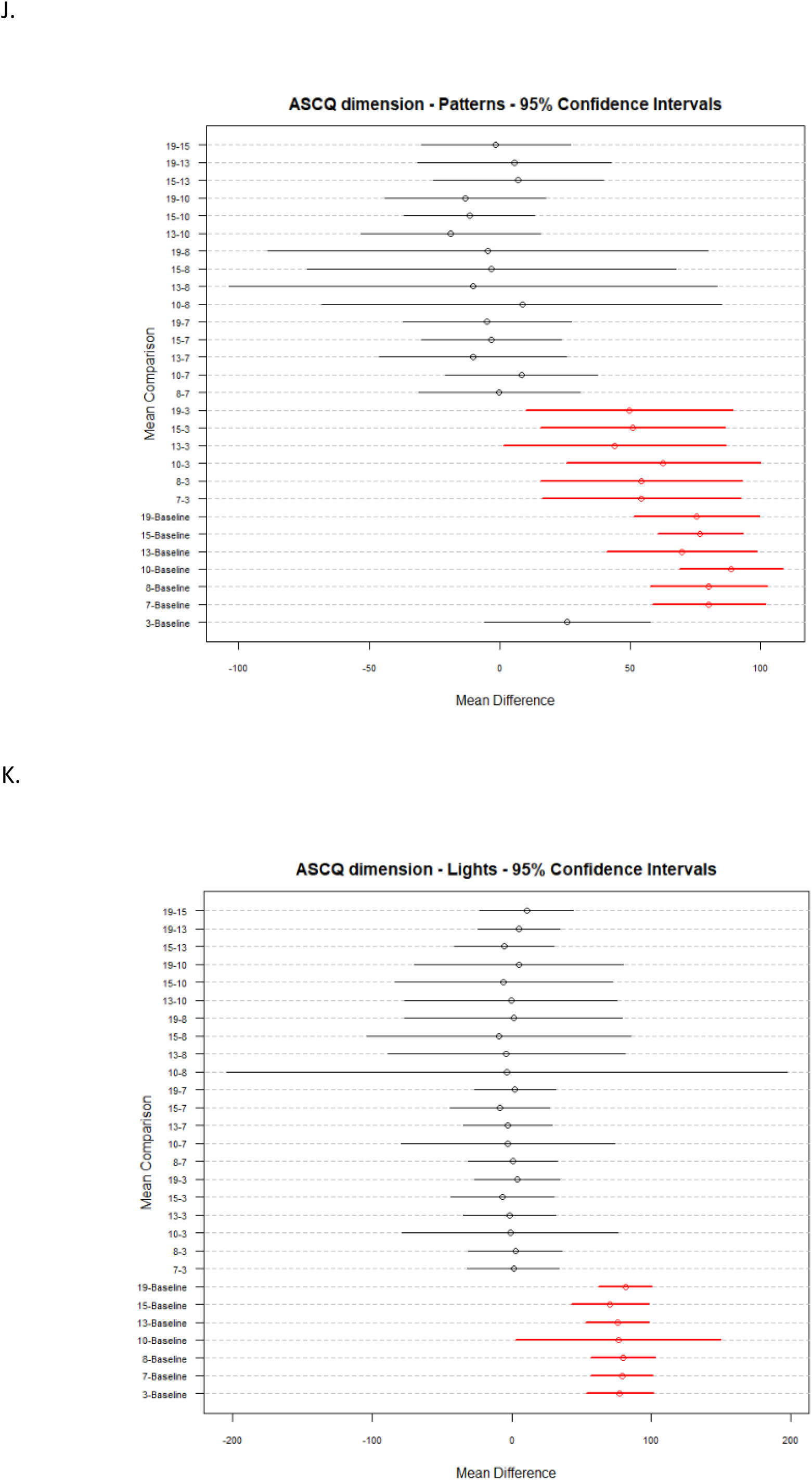
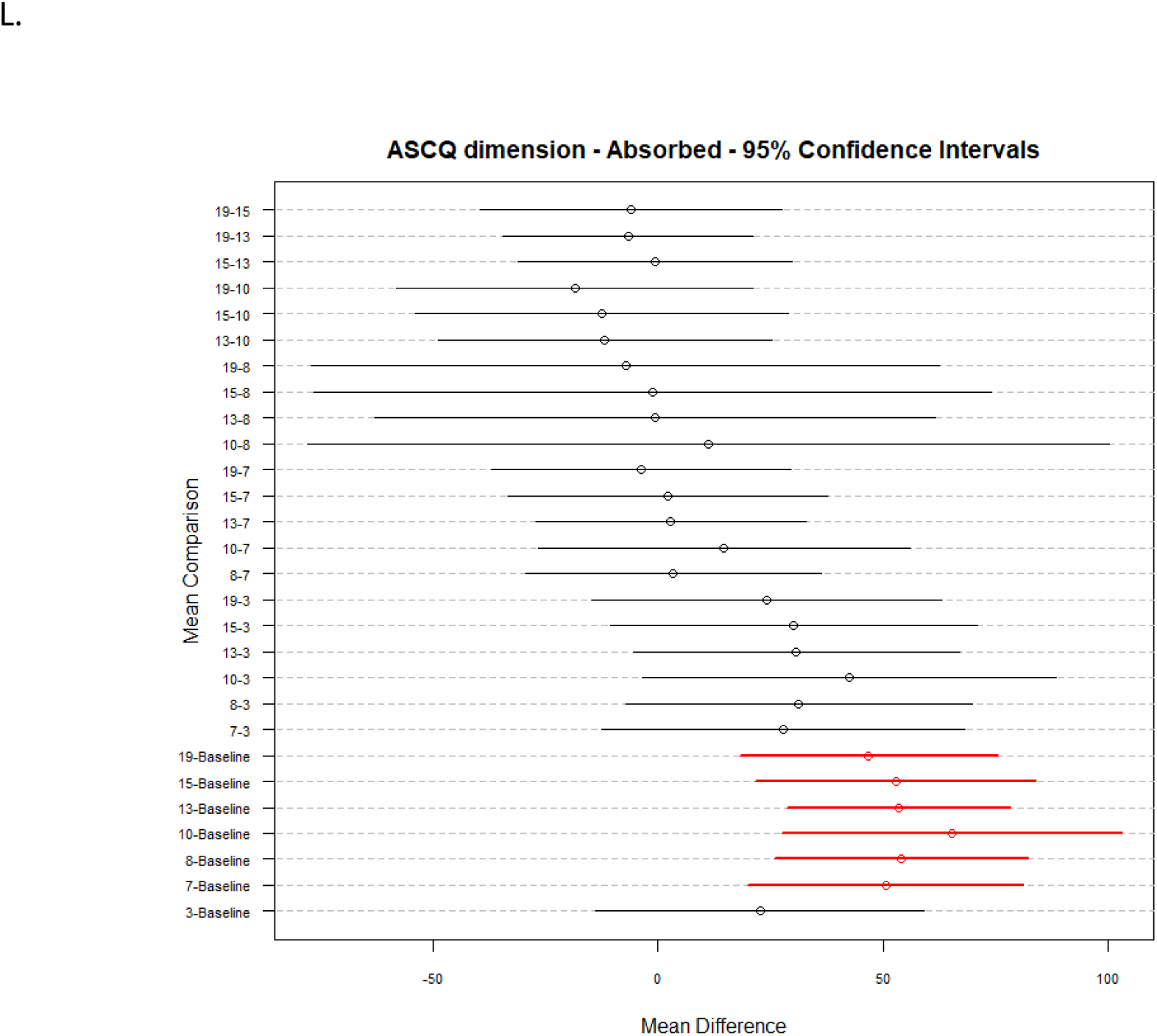
DTK Pairwise Multiple comparisons test based 95 % confidence intervals for ASCQ dimensions. The figures show the 95% percent confidence intervals for each of the pairwise comparisons between conditions for the following ASCQ dimensions Intensity (A), Imagination (B), Awe (C), Body (D), Colours (E), Surroundings (F), Float (G), Scene (H), Vivid (I), Patterns (J), Lights (K) and Absorbed (L) The significant pairwise comparisons are highlighted in red.

A Total measure was also calculated by adding the scores of all the other ASCQ dimensions and normalizing them across the number of dimensions (Fig 6A). A DTK test was also performed across this total measure for making comparisons between conditions (Fig 6B).

**Figure 6:**
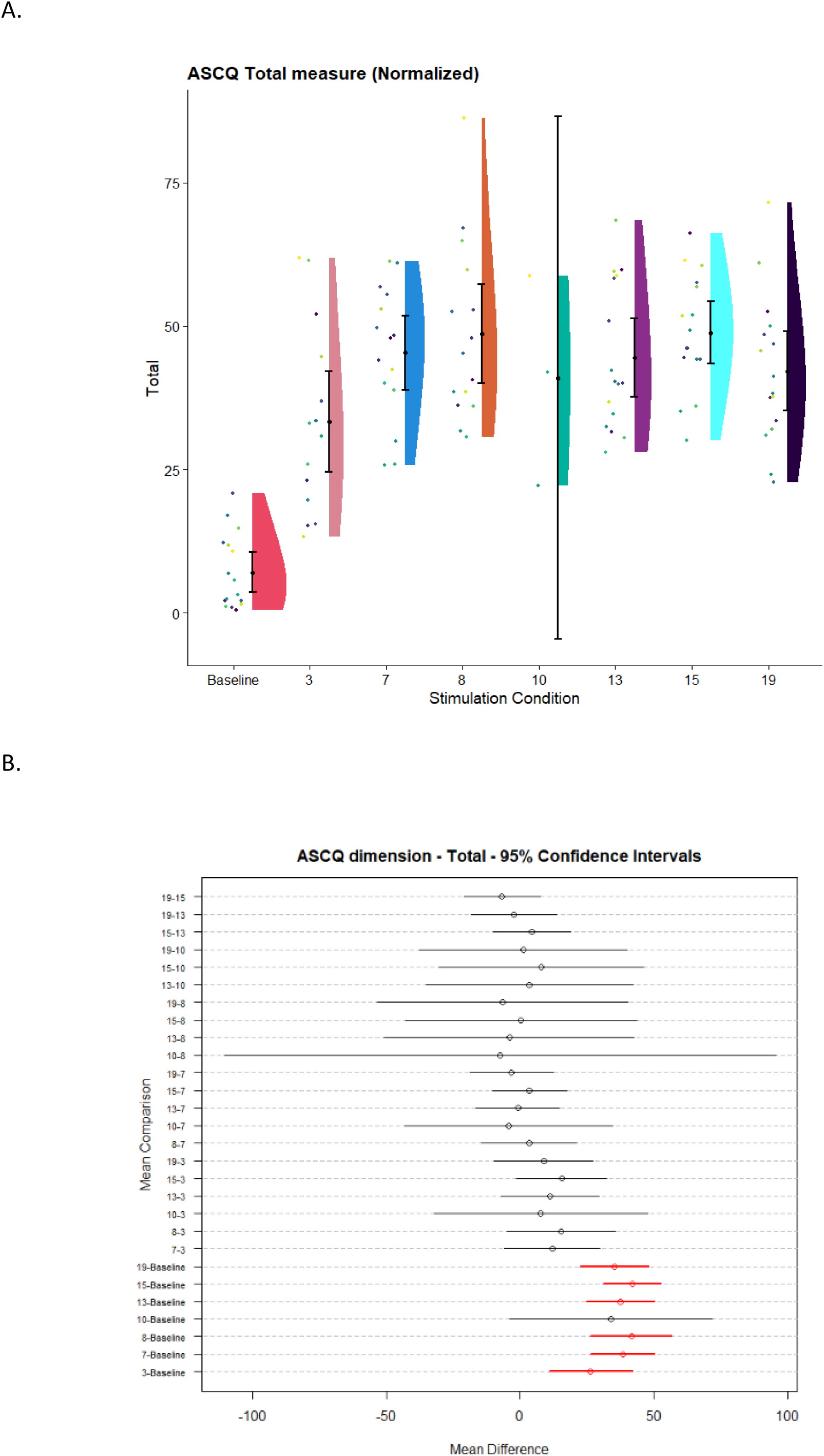
ASCQ total measure raincloud plot and DTK pairwise comparisons plot. Figure 6A shows the normalized ASCQ total scores across all conditions. Each participant’s values are represented with a uniquely coloured dot. The flat violin plots represent the distribution for each condition. The error bars represent the 95% confidence intervals of the data at each condition. Figure 6B shows the 95% percent confidence intervals for each of the pairwise comparisons between conditions for the total score with the significant interactions highlighted in red.

The results reveal a significant increase across most of the positive ASCQ dimensions, with stimulation conditions such as 8 Hz and 13 Hz being significantly different across all post-hoc tested dimensions except Imagination and Float when compared to baseline. Condition 8 Hz seemed to show the highest significant increase across dimensions Awe, Body, Surroundings, which was followed by 10Hz, 13 Hz and 7 Hz. These results seem to indicate that 8 Hz had the highest behavioural response, followed by 13 Hz. The 10 Hz condition seems to consistently have wide confidence intervals indicating a comparatively high level of variability in responses across participants for this condition; however, this might also be a result of the disproportionate number of participants for this condition as a result of the data loss. Similar to other measures, stimulation frequencies present within or close to the alpha band (8 - 12Hz) seemed to produce the highest increase all positive dimensions. The total measure indicated an overall increase, in comparison to baseline, up to 8 Hz after which it seemed to level off.

### Absolute spectral power

To compare changes in the spectral power profile across conditions, the EEG signal was decomposed into time, and frequency components for each condition and the absolute power across the spectral bands, delta (1 - 4 Hz), theta (4 – 8 Hz), alpha (8 – 13 Hz) and beta (13 - 30 Hz) was calculated and compared using paired t-tests with Bonferroni correction (Fig 7). For the theta band, there was a significant decrease across condition 3 Hz (t(13) = 4.146, *p_bon_* = .032) relative to the baseline dark condition. For the alpha band, there was a significant decrease across conditions 3 Hz (t(13) = 4.094, *p_bon_* = .036), 7 Hz (t(13) = 4.279, *p_bon_* = 0.025), 8 Hz (t(13) = 4.179, *p_bon_* = .030), 10 Hz (t(13) = 4.112, *p_bon_* = .034), 15 Hz (t(13) = 4.134, *p_bon_* = .033) relative to the baseline dark condition. This drop in alpha power is in line with past research with psychedelic and stroboscopically induced ASC (Rule et al., 2011; Michael M Schartner et al., 2017). There were no significant differences in beta or delta power across conditions. Figure 7 shows the distribution, 95% confidence intervals and the mean individual spectral power of each unique participant across the different conditions for each band.

**Fig 7:**
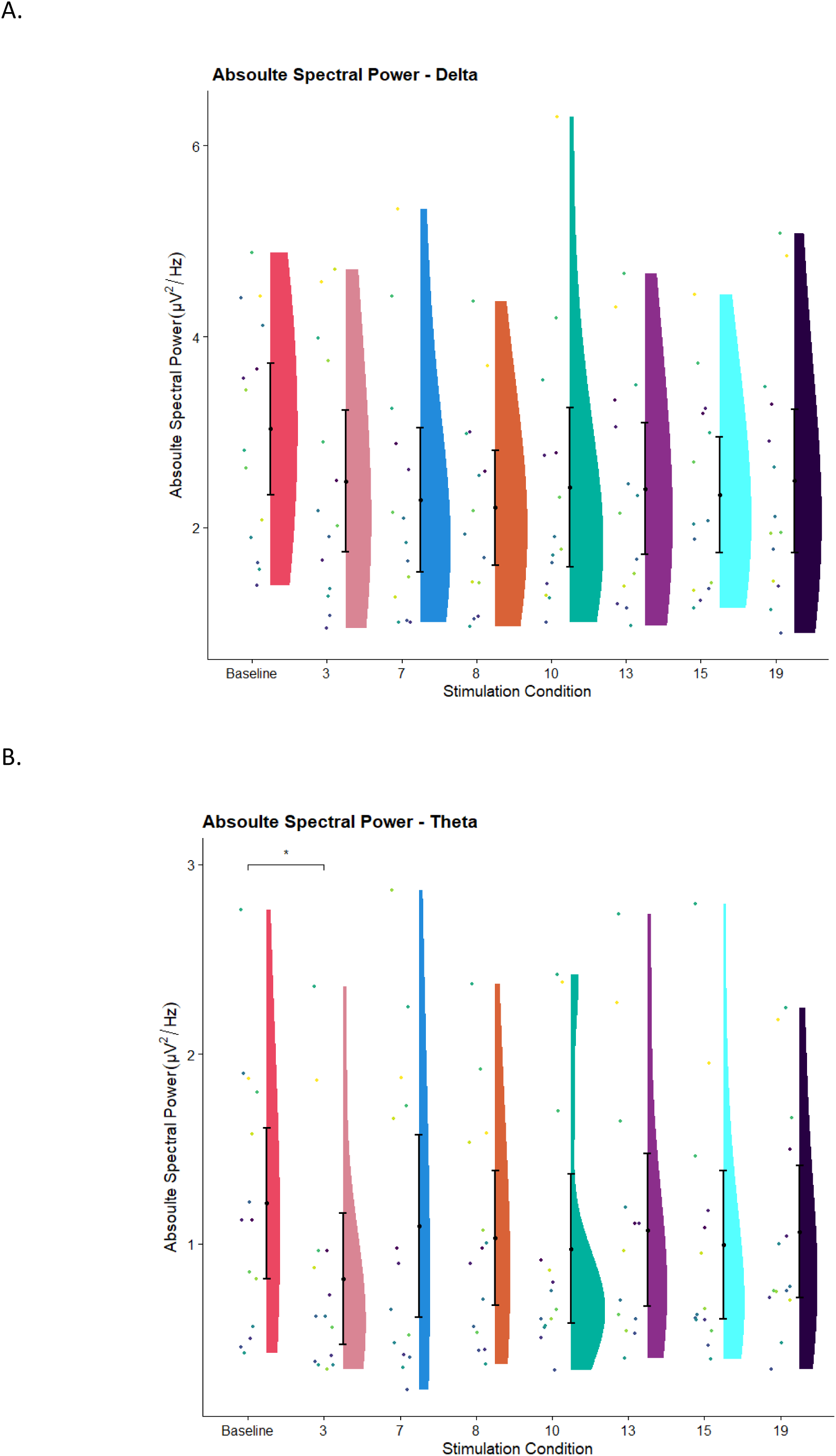
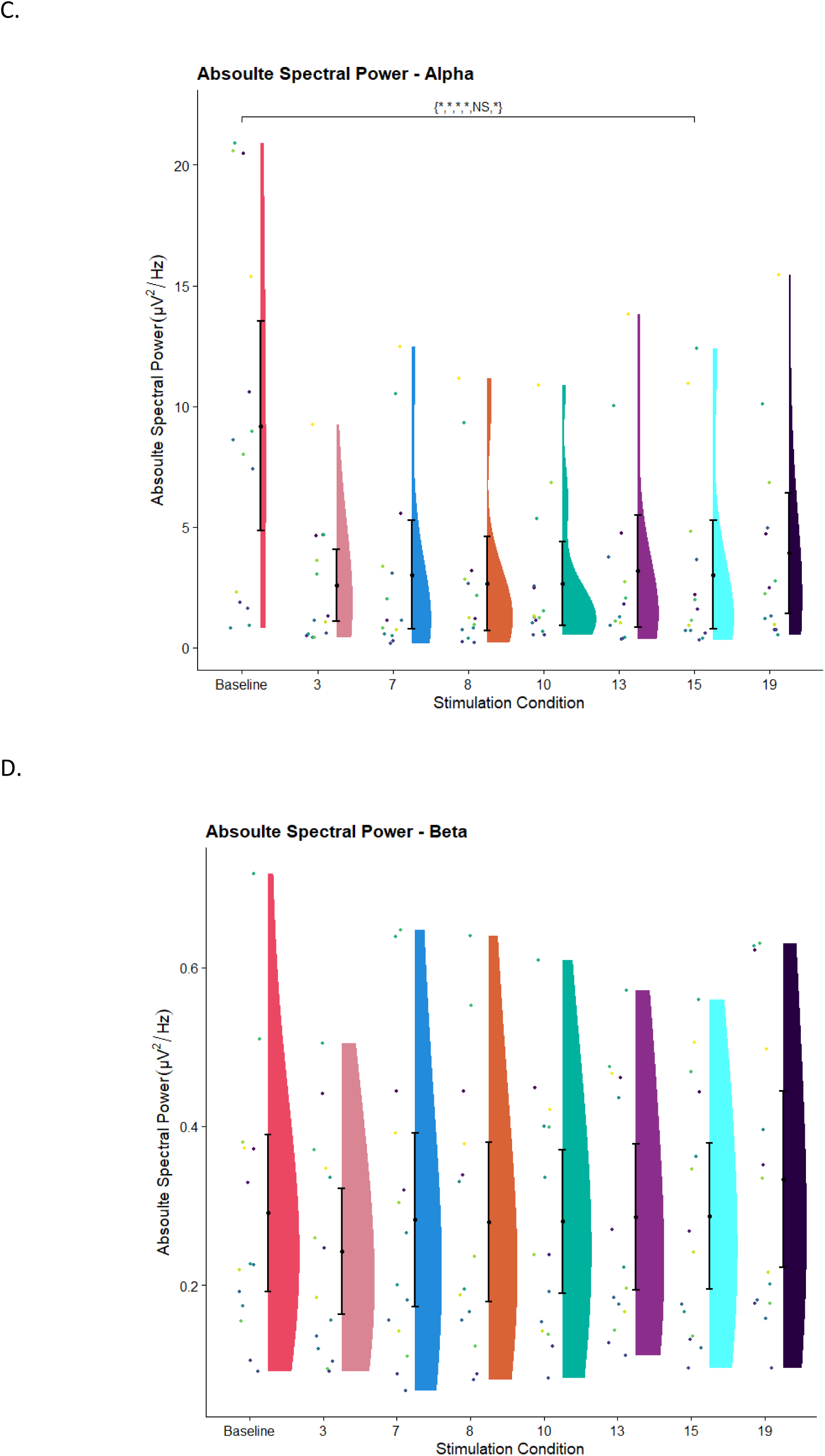
Absolute Spectral power. Rain cloud plots displaying the distribution of Delta (1 – 4 Hz), theta (4 – 8 Hz), alpha (8 - 12 Hz) and beta (13 – 30 Hz) power, averaged over a 4-minute window, across all conditions. Plot 7A shows the absolute delta spectral power, 7B shows the absolute theta spectral power, 7C shows the absolute alpha spectral power and 7D shows the absolute beta spectral power. Each participant’s values are represented with a uniquely coloured dot. All comparisons are Bonferroni corrected, and asterisks show significant comparisons, NS for non-significant, * indicates p < 0.05. The flat violin plots represent the distribution for each condition. The error bars represent the 95% confidence intervals of the data at each condition.

For theta power, 3 Hz was found to be lower than baseline; this result potentially reflects on how the stimulation frequency could alter the power across frequency bands close to the stimulation frequency.

### Normalized spectral power

Normalized spectral power was calculated so as to make comparisons with other studies using stroboscopic stimulation and psychedelics. To compare the changes in the spectral power density, the EEG signal was decomposed into time and frequency components, for each condition and the relative power across the spectral bands delta (1 - 4 Hz), theta (4 – 8 Hz),alpha (8 – 13 Hz) and beta (13 - 30 Hz) was calculated by normalizing the power across the four bands and compared using paired t-tests with Bonferroni correction. For the alpha band, there was a significant decrease in normalized power across all stimulation conditions, 3 Hz (t(13) = 6.559, *p_bon_* = < .001), 7Hz (t(13) = 6.229, *p_bon_* = < .001), 8 Hz (t(13) = 6.887, *p_bon_* = <.001), 10Hz (t(13) = 5.791, *p_bon_* = .002), 13 Hz (t(13) = 5.988, *p_bon_* = .001), 15Hz (t(13) = 6.142, *p_bon_* = < .001), 19 Hz (t(13) = 5.243, *p_bon_* = .004) respective to baseline dark condition. In addition, conditions 8Hz (t(13) = - 4.831, *p_bon_* = .009), and 15 Hz (t(13) = - 3.963, *p_bon_* = .045) were significantly lower than the 19 Hz condition. This drop in relative alpha power (in comparison to baseline) is in line with past research with psychedelics ASC and stroboscopic ASC (Michael M Schartner et al., 2017; Schwartzman et al., 2019). For the beta band, there was a significant increase in spectral density across all stimulation conditions, 3 Hz (t(14) = - 6.161, *p_bon_* = < .001), 7 Hz (t(14) = - 7.567, *p_bon_* = < .001), 8 Hz (t(14) = - 7.279, *p_bon_* = < .001), 10 Hz (t(14) = - 7.696, *p_bon_* = < .001), 13 Hz (t(14) = - 7.863, *p_bon_* = < .001), 15 Hz (t(14) = - 7.114, *p_bon_* = < .001), 19 Hz (t(14) = - 7.189, *p_bon_* = < .001) respective to the baseline dark condition. There were no significant differences in theta power spectral density and delta power spectral density across conditions. Figure 8 shows the distribution, 95% confidence intervals and the mean individual spectral power of each participant across the different conditions for each frequency band.

**Fig 8:**
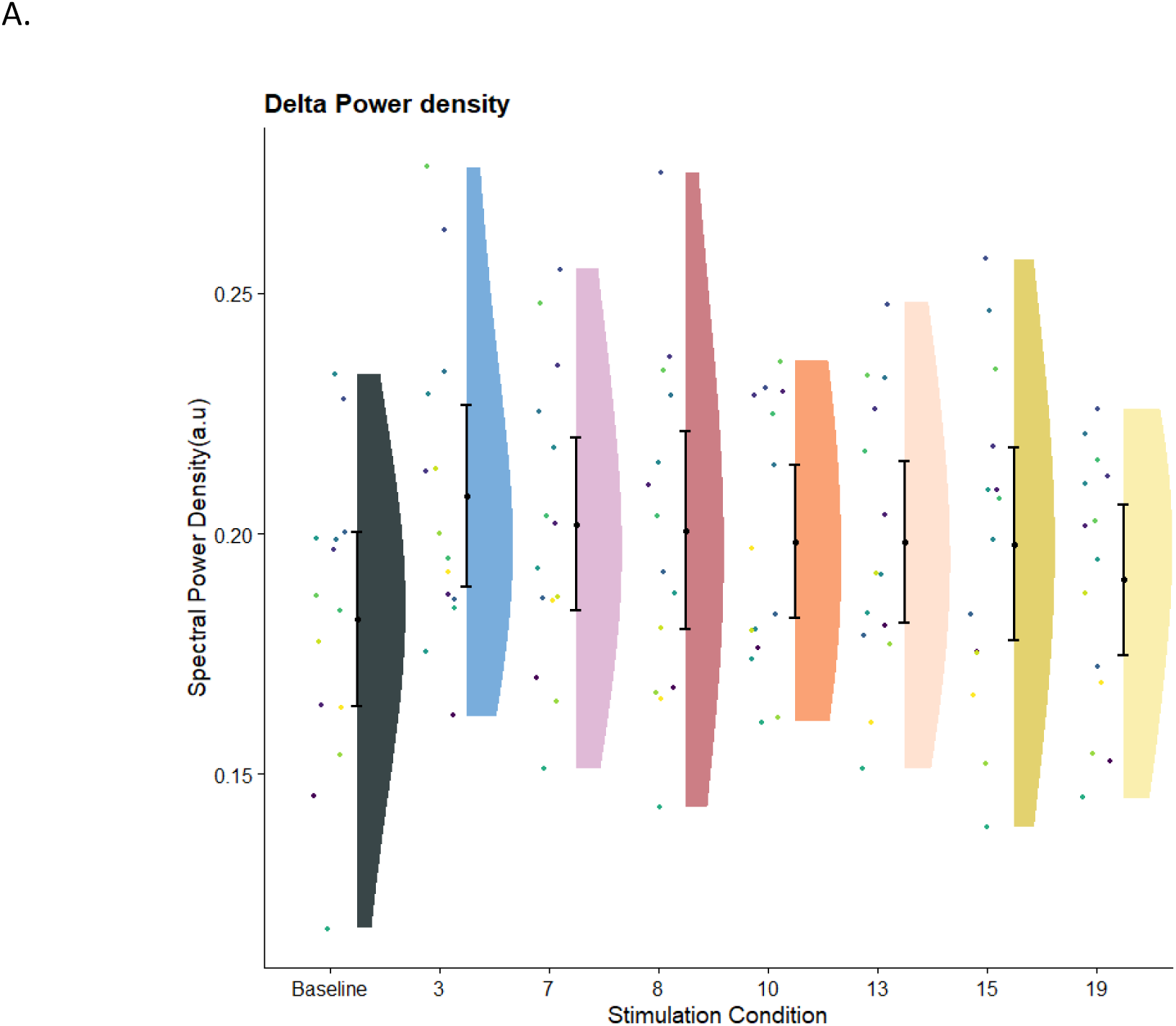
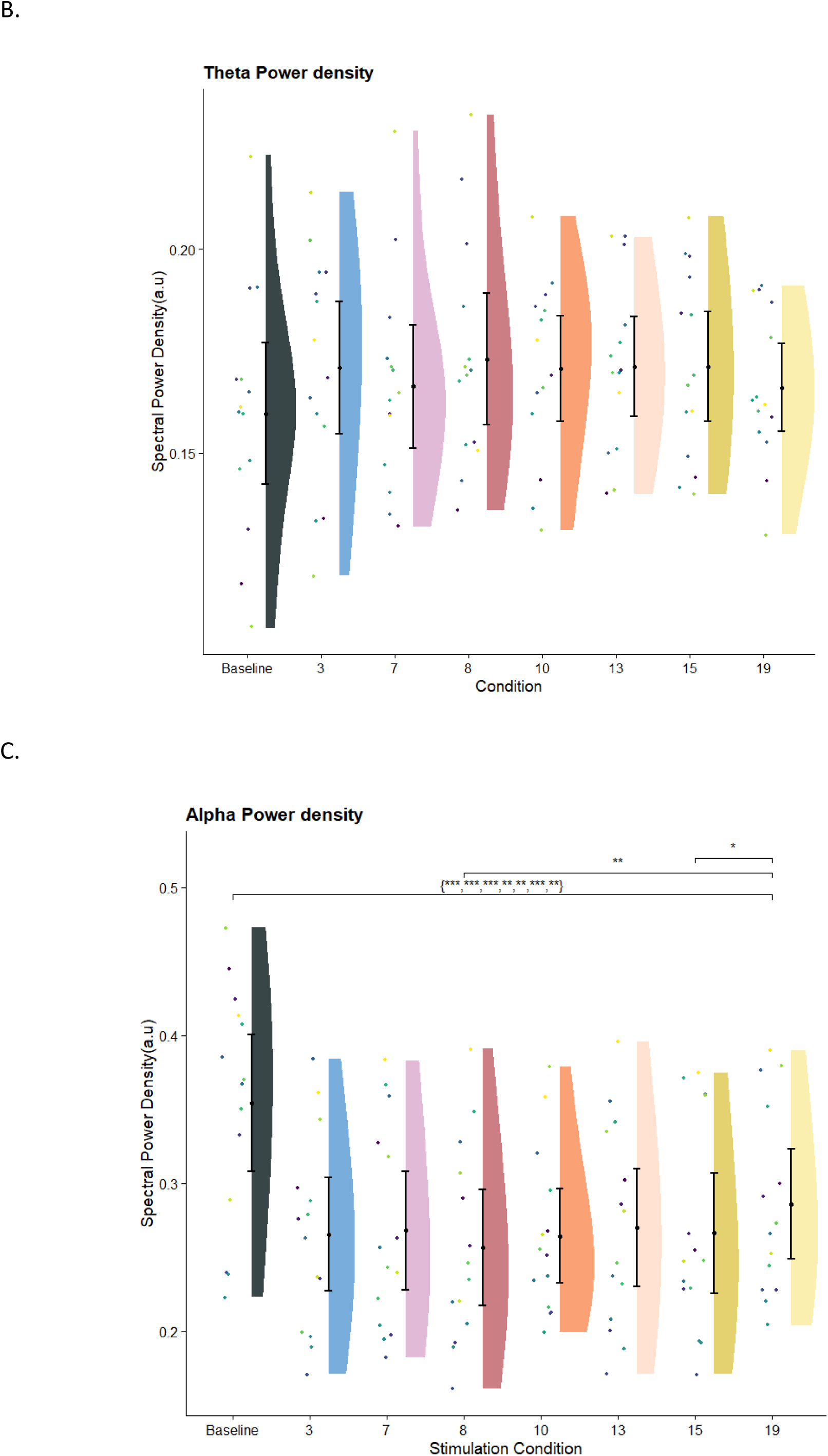
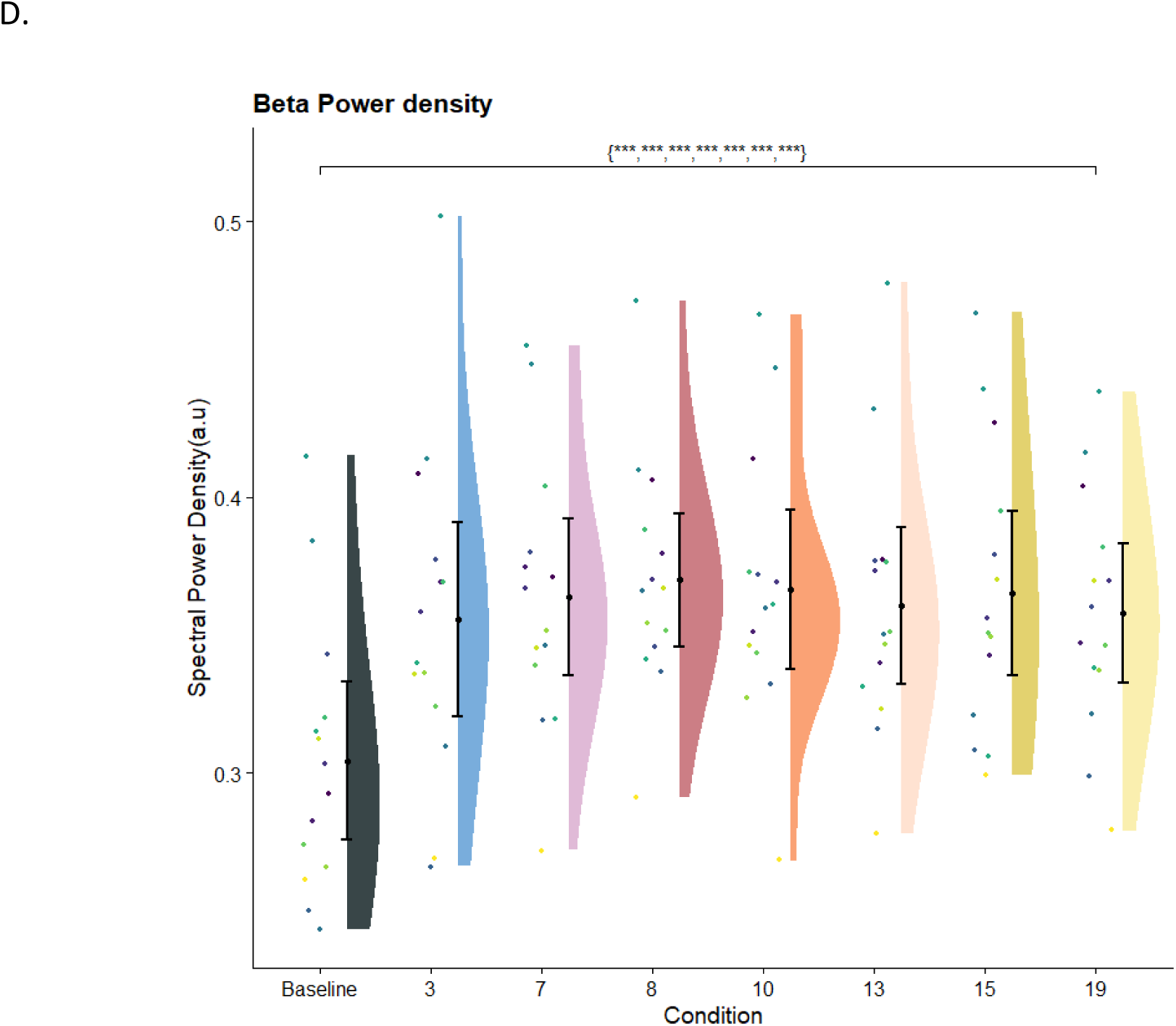
Normalized spectral power. The plot displays the distribution of Delta (1 – 4 Hz), theta (4 – 8 Hz), alpha (8 - 12 Hz) and beta (13 – 30 Hz) power spectral density, averaged over a 4-minute window, for all participants across different conditions. Plot 8A shows the delta power spectral density, Plot 8B shows the theta power spectral density, Plot 8C shows the alpha power spectral density and Plot 8D shows the beta power spectral density. All comparisons are Bonferroni corrected and significant comparisons are shown by asterisks, NS for non-significant, * indicates p < 0.05, ** indicates p < 0.01, *** indicates p < 0.001 and **** indicates p < 0.0001. The flat violin plots represent the shape of the distribution for each condition. The individual coloured dots represent individual participants results with each participant represented with a unique colour. The error bars represent the 95% confidence intervals of the data at each condition.

In the alpha power density plot (Fig 8C), there seems to be a clear increasing trend in spectral density across stimulation conditions, with higher alpha spectral density for higher stimulation frequencies. This result could be reflective of the associated increase in the level of visual activity as the stimulation frequency increases. The beta power density also seems to have increased across all stimulation conditions with 8 Hz showing the maximal effect across conditions.

### Correlation analysis between signal diversity, power spectra, spectral density and ASCQ dimensions

In order to investigate possible relationships between neural measures and subjective reports, we calculated the relative difference between each stimulation condition versus their baseline scores, i.e. stimulation condition - baseline for signal diversity (LZs), absolute power spectral (Alpha, Beta, Delta and Theta), spectral density (Alpha, Beta, Delta and Theta) and each ASCQ dimension. We then examined the correlations between them using Pearson correlations for each stimulation condition (Fig 9). Figure 9 only displays correlations, where p < .01. Given the high number of variables, the correlations were not corrected for multiple comparisons.

**Figure 9:**
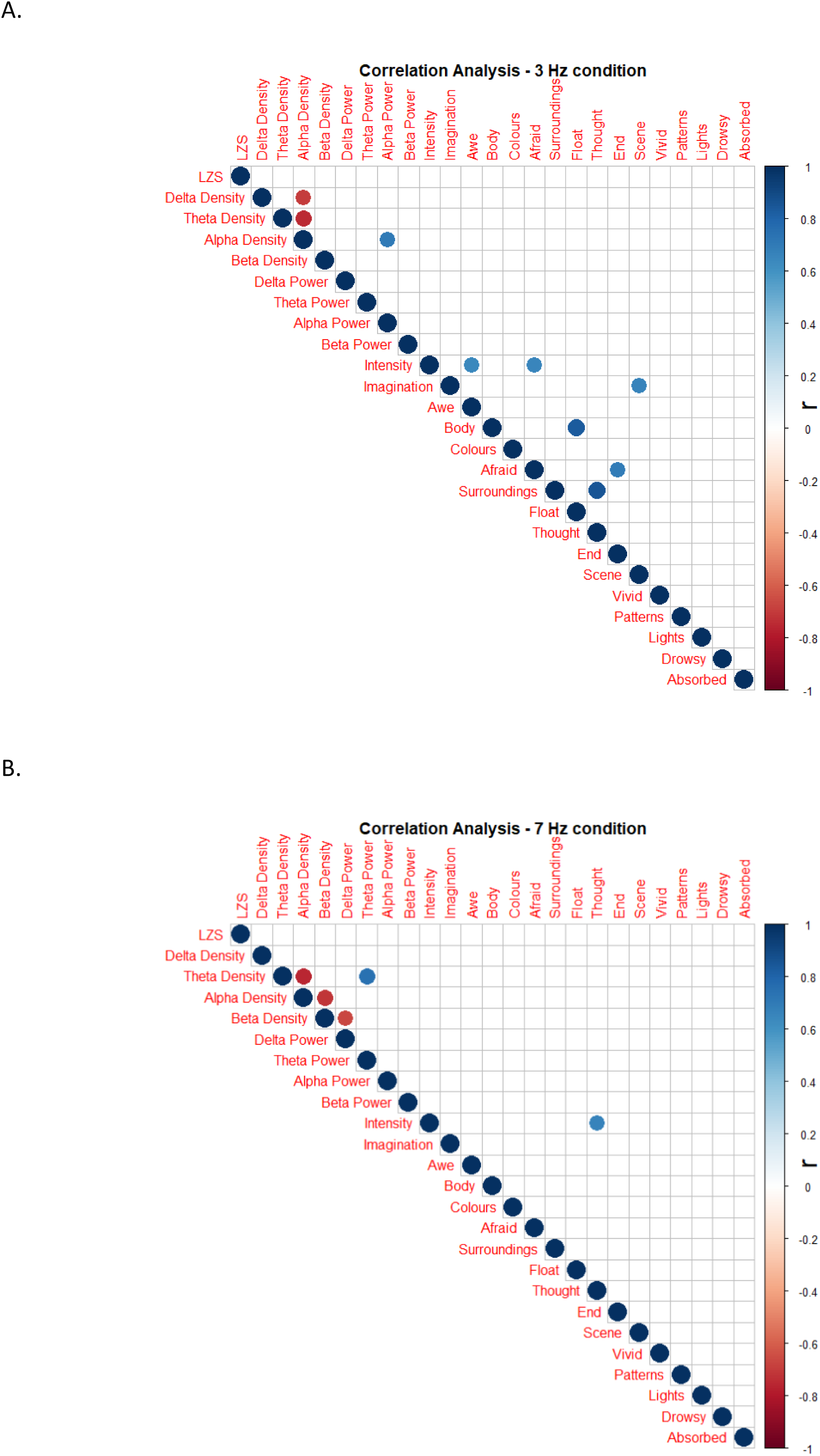
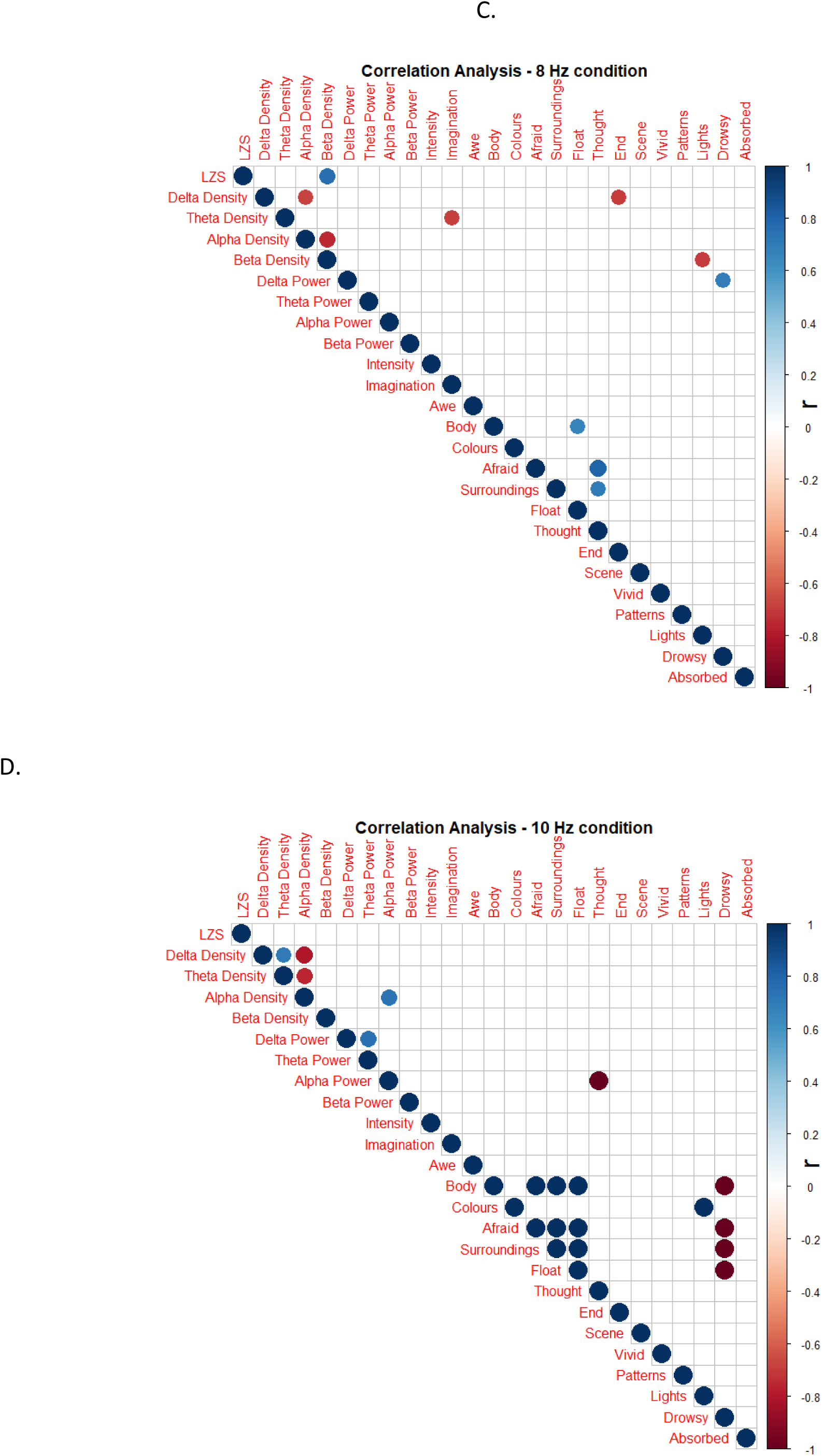
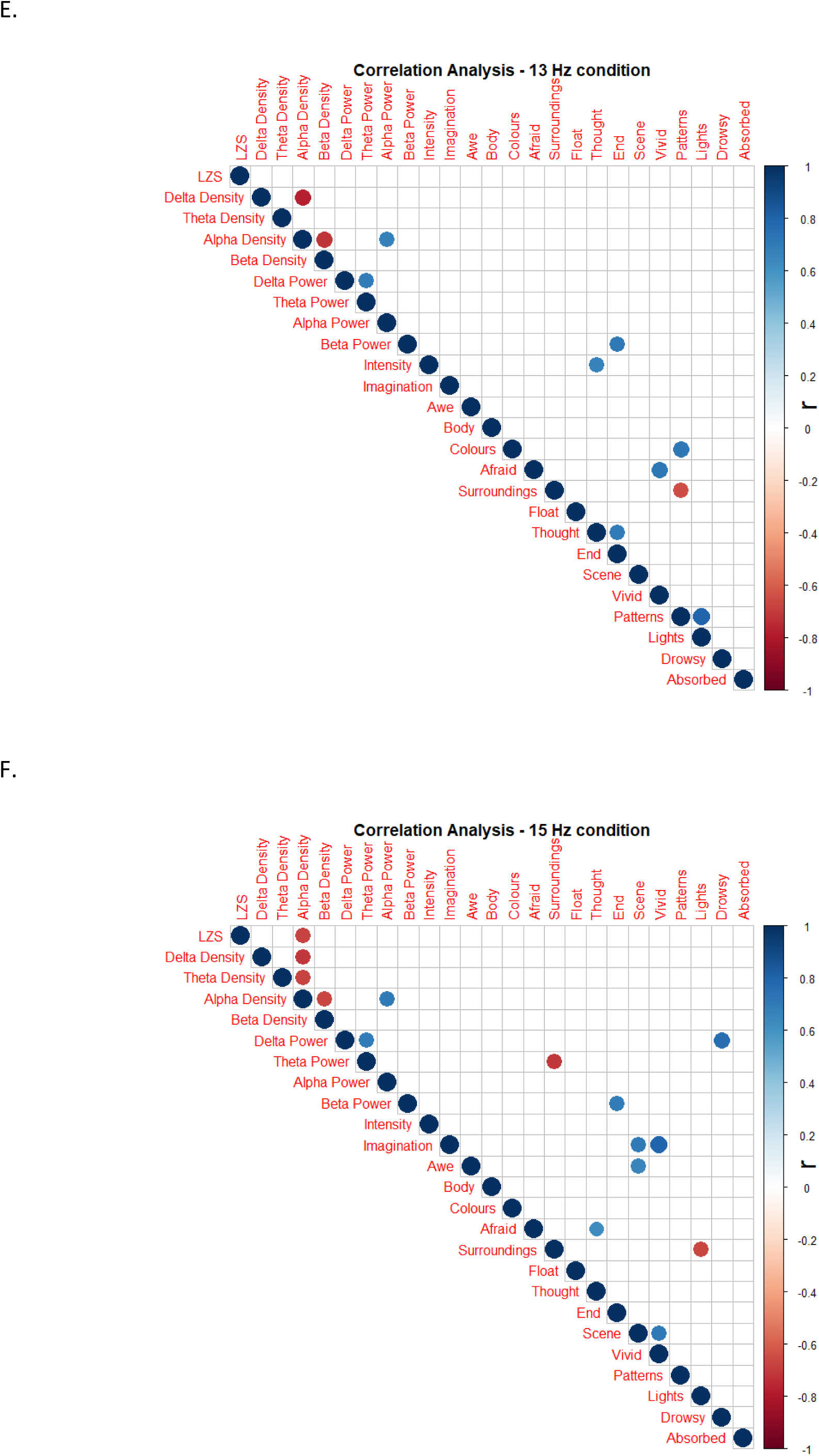
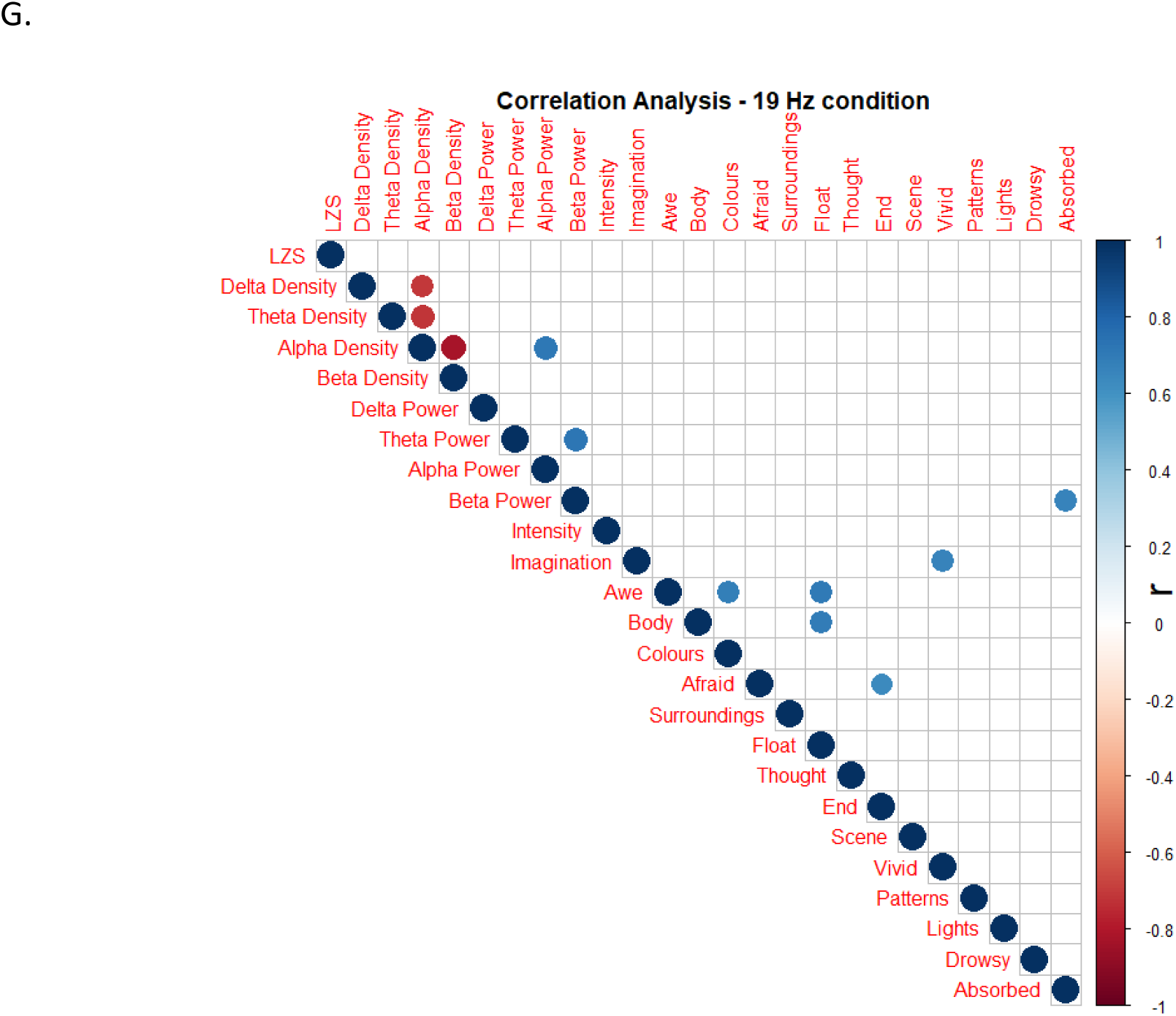
Correlation matrix showcasing Pearson correlations of relative difference across neural measures and questionnaire dimensions for all stimulation conditions. For each condition, a matrix indicates the Pearson correlation(r), of the score difference between each stimulation condition and the baseline dark condition. Plot 9A shows the correlation matrix for condition 3 Hz, Plot 9B shows the correlation matrix for condition 7 Hz, Plot 9C shows the correlation matrix for condition 8 Hz, Plot 9D shows the correlation matrix for condition 10 Hz, Plot 9E shows the correlation matrix for condition 13 Hz, Plot 9F shows the correlation matrix for condition 15 Hz, Plot 9G shows the correlation matrix for condition 19Hz. The matrices have blank white squares where the significance value (p) is below 0.01 or if r = 0. Since these correlations are exploratory, they have not been controlled for family-wise errors. The correlation values are represented from Dark Blue for r = 1 and range up to Dark Red for r = −1.

Alpha density was found to be strongly correlated with alpha power across all stimulation conditions except 7 Hz and 8 Hz, which is probably indicative of the drop in alpha spectral power and density. Alpha density was found to have a strong inverse correlation with delta density for all conditions except 7Hz and with theta density for all conditions except 8 Hz and 13 Hz. Particularly within the 8 Hz condition, delta density has a strong inverse correlation with the ASCQ dimension End. Unsurprisingly, delta power had a strong correlation with Drowsy, and Lights had a strong inverse correlation with Beta density, Thought had a strong correlation with Afraid and Surroundings. Within the 10 Hz stimulation condition, Body, Afraid, Surroundings and Float all seem to be very strongly intercorrelated and drowsy being strongly inversely correlated with them.

## Discussion

The study aimed to explore if there was any relationship between the stimulation frequency and different measures of neural dynamics, and the difference in experience of stroboscopically induced visuals. As expected, the stroboscopic stimulation produced visual stimulations across all stimulation frequencies. The level of subjective changes produced by the stroboscopic stimulation, increased in the order of increasing frequency, with the weakest overall experience being reported at 3 Hz and the most intense experience at 19 Hz, as indicated by the average scores across the ASCQ questionnaire.

In terms of the intensity of the experience, the 8 Hz condition had the highest scores followed by 7 Hz, 19Hz, 15Hz, 10 Hz, 13 Hz and 3 Hz. In terms of the reported visual characteristics, condition 10 Hz had the highest difference from baseline as indicated by the results of dimensions Colors, Patterns and Absorbed, followed by 8 Hz, 13Hz and 7 Hz. The ASCQ total scores also indicated a similar picture with 8 Hz and 15 Hz producing the highest scores, with an increase in experience across all conditions.

### Spectral Power profile changes

Within the spectral power profile, the absolute power results indicated a general decrease in alpha power across all stimulation conditions and a general decrease in theta power with a maximal decrease for the 3 Hz condition. The power spectral density results (relative spectral power) indicated an increase in beta power density across all stimulation conditions and a general decrease in alpha power density across all conditions.

### Signal Diversity

One of the key findings of the study is the variation in signal diversity across different stimulation conditions, which was similar to psychedelic induced ASC’s with increased signal diversity for all conditions. As expected, all conditions displayed higher LZ scores than baseline dark condition, with the 8 Hz condition showing the maximal increase in LZs, followed by 10 Hz and 7Hz. The results seem to show an increasing LZs score till the 8 Hz condition, after which LZs scores levelled off. This result seems to indicate that all the frequencies tested seem to produce ASC’s similar to that of psychedelics and that signal diversity can effectively measure such phenomenological differences in conscious content.

These results also highlight the importance of the alpha band stimulation frequencies in creating the most intense stroboscopic experiences and also the maximal changes across the ASCQ and the spectral profile in this study, lending credence to the use of such frequencies for stroboscopic stimulation for maximum effect in future studies. Based on the ASCQ results, the 8 Hz stimulation frequency produced the most pronounced consistent level of changes among the frequencies tested. The 10 Hz, 7 Hz and 13 Hz frequencies produced similarly high results across the ASCQ, but with participants having a clearly higher response for one frequency over another rather following any linear patten. As suggested by Bartossek et al. (2020), this level of variability might depend upon on the participant’s individual alpha rhythm and matching stimulation frequency to individual alpha in the future may result in a much more intense experience (Koch, Steinbrink, Villringer, & Obrig, 2006). Higher frequencies such as 19 Hz and 15 Hz also produced similarly high ASCQ results, but had higher ratings on the dimension Scene, which might indicate a potentially more complex SIVH compared to other frequencies.

Spectrally, all stimulation conditions displayed a significant decrease in alpha power and alpha power density, which is similar spectral response to those induced by psychedelics and has been considered by some models as markers of the underlying psychedelic state (Riba et al., 2004; Rule et al., 2011; Michael M. Schartner et al., 2017). Past research has indicated that the decrease in alpha power is potentially due to the broadband cortical desynchronization that occurs during psychedelic state across several regions of the brain (Robin L. Carhart-Harris et al., 2012; Muthukumaraswamy et al., 2013). This desynchronization of the brain activity might potentially disrupt the top-down and bottom-up processes relating to predictive processing accounts of perception, thereby creating such an ASC (R. L. Carhart-Harris & Friston, 2019; Robin Lester Carhart-Harris et al., 2014; Friston, 2010). The results seem to indicate that this shift in alpha power might be a correlate of the altered states of consciousness and the underlying human psychedelic state and that psychedelics might induce such neural entrainment through pharmacological interactions.

A consistent feature across all tested measures was the level of variability in results shown by participants for all conditions. While the general trends were still retained across conditions, certain participants exhibited a lot wider range (up to 3 times greater between conditions) of spectral power responses to a specific stimulation condition while being closer to the median at other stimulation frequencies. Given past research with psychedelic ASC’s, the level of individual difference may depend on the physical and structural differences of the early visual system cells such as hypercolumns and retinal ganglion cells (Rule et al., 2011), or the level of associated neurotransmitter release and activity (2NMA and 5HTP) (Muthukumaraswamy et al., 2013). This level of individual difference has not been observed in psychedelic ASC’s and might play an essential role in understanding how brain synchronizations occurs and potentially even in understanding the underlying hallucinatory state associated with such ASC’s. Even across a particular stimulation frequency, the type of visual effects seen may vary across participants and could even have multiple such stimuli and form constants occur in a multi-stable form (Allefeld, Pütz, Kastner, & Wackermann, 2011). Future research is needed to explore the underlying mechanism that account for the individual differences in neural responses to stroboscopic stimulation.

### Limitations

One of the critical limitations of this study is that while several stimulation frequencies were tested there were still frequencies within these bands that were not tested and did not account for minor frequency changes (< 1 Hz). Other frequencies that were not tested in this study might showcase a different pattern of results even within the same spectral bands. A significant limitation to note with signal diversity measures is that while they have been showcased to be reflective of conscious content and level, they might not encompass all facets of consciousness and should not be taken as an indicator of consciousness as a whole (A. K. Seth et al., 2006). Future studies should focus on using a dynamic frequency sweep across all the frequencies to map the responses completely.

### Conclusion

The study shows the breadth of neural differences possible with stroboscopic stimulation using different stimulation frequencies. Alpha band (8 – 12 Hz) and close to alpha band frequencies seemed to produce the highest-level changes across the spectral profile, signal diversity and in subjective reports. Of the alpha band frequencies tested, 8 Hz consistently produced the maximal change across neural measures and seemed to have the highest shift in subjective report relative to baseline. The level of variability across these different conditions varied significantly for each participant, with some participants being quite responsive to one frequency while being median at another. This variability might be a result of variations in the underlying natural rhythms of the brain, and more research is required to explore this idea further. Overall, the study shows the level of specificity of modulation possible with stroboscopic stimulation and its potential in the research of ASC’s without the use of psychedelics.

## Acknowledgement

I would like to thank my supervisor, Dr David Schwartzman, for the consistent support, encouragement and pushing me further. Without his guidance and support, this dissertation would have never been possible.

In terms of the original study, it was designed and conducted by Dr David Schwartzman, and Ales Oblak and the author helped with data collection. All the data processing and statistical analysis were prepared and performed by the author with input from Dr Schwartzman. Thanks to Dr Lionel Barnett for the signal detrend script and to Dr Anil Seth and The Sackler Lab for providing the opportunity for me to be a part of this study.

## Notes

### Competing Interest Statement

The authors have declared no competing interest.

