## Appendix for "Neural dynamics of stroboscopic stimulation at different stimulation frequencies"

Appendix A

Lempel-Ziv Complexity across all channels (LZc)

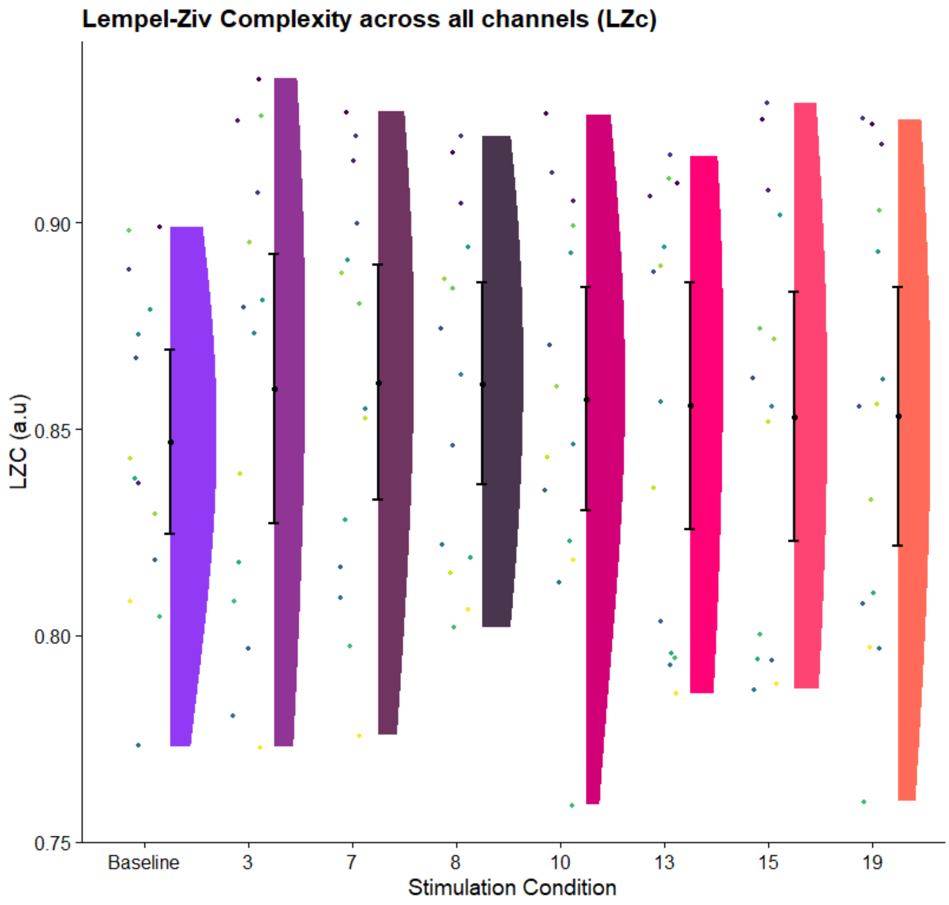


Figure: Mean Lempel-Ziv complexity scores (LZC). Rain cloud plot displays the Lempel-Ziv complexity scores over all channels, averaged over each conditions 4-minuite window, and all participants. All comparisons are Bonferroni corrected and significant comparisons are denoted by asterisks, NS for non-significant, * indicates $p < 0.05$. The violin plots represent the distribution for each condition. The individual coloured dots represent individual participants results with each participant represented by a unique colour. The error bars represent the 95% confidence intervals of the data for each condition.

Appendix B

Altered States of Consciousness Questionnaire (ASCQ)

1. 'Please rate the intensity of the experience during the last session',
2. 'I could see pictures from my past or my imagination extremely clearly '
3. 'I experienced a sense of awe'
4. 'It seemed to me as if I didn’t have a body or a body part anymore'
5. I saw colours before me'
6. 'I was afraid without being able to say exactly why'
7. 'I experienced my surroundings as being strange and weird'
8. 'I felt as though I was floating'
9. 'I was not able to complete a thought, my thought repeatedly became disconnected'
10. 'I wanted this state to end as soon as possible'
11. 'I experienced the scene rolling by my mind’s eye'
12. 'My imagination was extremely vivid'
13. 'I saw regular patterns'
14. 'I saw lights or flashes of lights in my mind’s eye'
15. 'I felt drowsy'
16. I was absorbed in the experience.
